# Malnourishment and expanded microbiome maintain superior efficacy of the recombinant tuberculosis vaccine BCG::ESAT-6-PE25SS

**DOI:** 10.64898/2026.08.16.743391

**Authors:** Munish Puri, Manoharan Kumar, Haleagrahara Nagaraja, Harindra Sathkumara, Serrin Rowarth, Kylie Robertson, Selvakumar Subbian, Jeffrey Warner, Catherine Rush, Roland Ruscher, Matt Field, Andreas Kupz

## Abstract

Tuberculosis (TB) remains the leading cause of infectious mortality. The limited efficacy of the only TB vaccine, Bacille Calmette–Guérin (BCG) against pulmonary disease necessitates improved vaccines. Host factors such as malnutrition and microbiome composition shape immune responses in humans, though these factors are overlooked in preclinical vaccine evaluation. Here we show that a recombinant BCG strain, BCG::ESAT-6-PE25SS, confers superior protection compared to BCG across murine models of malnutrition, antibiotic-induced dysbiosis and environmentally enriched microbiota. Unexpectedly, malnourished mice displayed reduced *Mycobacterium tuberculosis (Mtb)* burden, associated with altered host metabolism and immune composition. Microbiome disruption increased TB susceptibility, whereas diversification of microbiota enhanced resistance and immune heterogeneity. Vaccine efficacy correlated with enrichment of known immunomodulatory microbial taxa. These findings suggest diet-microbiome-immunity interactions as potential key determinants of TB pathogenesis and provide evidence for the importance of vaccine candidate evaluation under physiologically relevant co-morbid conditions

## Introduction

Tuberculosis (TB) remains the leading cause of infectious death worldwide. In 2024, an estimated 10.7 million people contracted TB, and 1.23 million died from the disease globally^1^. In low- and middle-income countries pre-existing communicable and non-communicable immunomodulatory conditions are common and contribute to TB incidence and progression of *Mycobacterium tuberculosis (Mtb)* infection^2, 3^. In particular, the association between TB and comorbidities such as HIV^4–6^ and diabetes^7–12^ has been extensively studied. Both conditions significantly increase host susceptibility to TB. In addition, a wide range of socioeconomic determinants, including poverty, overcrowding, and limited access to healthcare further elevate TB risk^13–18^. Among biological risk factors, malnutrition is considered a major contributor to TB incidence and disease progression, primarily due to its detrimental effects on the immune system^19–22^. The World Health Organization (WHO) defines malnutrition as a deficiency, excess, or imbalance in a person’s intake of energy and nutrients. Malnutrition therefore includes both undernutrition and overnutrition, as well as deficiencies in specific vitamins or minerals. Clinically, severe undernutrition may present as kwashiorkor, which is commonly associated with oedema and protein deficiency, or marasmus, which is characterised by marked wasting due to prolonged inadequate energy intake^23^.

Bacille Calmette-Guérin (BCG) is the only licensed prophylactic TB vaccine, but its efficacy diminishes over time. While it provides strong protection against disseminated TB in children who have never been exposed to mycobacteria, its efficacy in adults is variable (0-80%) and differs among populations around the globe. Furthermore, BCG vaccination does not protect against TB in adults, who contribute significantly to disease transmission^24–26^. Therefore, more effective TB vaccines are urgently required to reduce the high morbidity and mortality associated with the disease.

While several TB vaccine candidates have not shown superiority over BCG in advanced clinical trials^4, 27, 28^, there are multiple TB vaccine candidates currently in pre-clinical or clinical trials, aiming to improve BCG-mediated immunity^29^. One such live-attenuated vaccine candidate is BCG::ESAT-6-PE25SS (abbreviated to PE25SS in Figures), a recombinant BCG strain (rBCG) that secretes full-length ESAT-6, an immunodominant secreted protein from *Mycobacterium tuberculosis* (*Mtb*) using the endogenous ESX-5 secretion system of BCG^30^. Although many TB vaccine candidates, including BCG::ESAT-6-PE25SS, have been evaluated in small animal models of TB, its efficacy against TB in comorbid immunomodulatory conditions remain unknown. It is increasingly recognised that the efficacy and immunogenicity of a vaccine depend on multiple factors, including host-associated factors such as nutrition, age, comorbidities and microbial diversity^31^.

Humans host trillions of commensal bacteria (microbiota) on external surfaces (e.g., skin) and internal organs (e.g., lungs and gastrointestinal tract). These microbes and their metabolic products, collectively referred to as the microbiome, play a vital role in immune regulation and homeostasis. Disruption of the microbiota, often due to antibiotic use, has been linked to the development of various diseases, including chronic respiratory diseases^32–36^. Antibiotic overuse, which is common in TB-endemic countries, tends to increase TB susceptibility and the emergence of multi drug-resistant TB cases.

Importantly, antibiotic-induced microbial dysbiosis in humans can be modelled and studied using antibiotic-treated animals^37–39^. Standard laboratory mice, typically raised in controlled, pathogen-free environments, are often referred to as ‘clean mice’. Their microbiota closely resembles that of human infants due to minimal environmental exposure^40, 41^. However, recent studies have introduced a more representative model involving co-housing laboratory mice with pet-store (or ‘dirty’) mice for 12–15 weeks. This exposure to diverse environmental microbes results in a more mature, human-like microbiota^40–42^. These ‘dirty mice’ models are now regarded as valuable and more predictive tools for immunological studies and vaccine testing^40^.

Here we assessed the role of diet and microbiota alteration on TB susceptibility and the efficacy of the BCG and BCG::ESAT-6-PE25SS vaccines in these comorbidities. Our results show that mice maintained on a customised energy-restricted, low-protein, low-fat, high-carbohydrate, high-fibre and ‘dirty’ mice are overall less susceptible to TB disease and that the BCG::ESAT-6-PE25SS vaccine outperforms BCG efficacy in malnourished and antibiotic-treated mice. In line with variability observed in human population studies^43–46^ ‘dirty’ mice also show a diverse response to TB vaccination.

## Results

### Malnourished mice exhibit attributes of human malnourishment

To study vaccine performance in malnourished conditions, we first established a mouse model of human marasmus. To this end, BALB/c mice were fed a fixed normal diet (ND) or malnourished diet (MD) diet of 30 g per cage for two weeks (**Fig. 1a).** During this dietary intervention, body weight was measured daily, and body length was recorded weekly. The malnourished mice (**Fig. 1b**, right) showed stunted growth compared to the ND-fed mice, with marked reductions in both body length (**Fig. 1c**), body weight (**Fig. 1d**) and body mass index (**Fig. 1e**). Malnourished mice consumed more food than SPF mice (**Fig. 1f)** and had significantly heavier faecal pellets than ND-fed mice (**Fig. 1g).** Additionally, faecal pellet colour differed between groups (ND: brown; MD: light green), and MD-fed mice organ weights, such as those of the liver (**Fig. 1h**) and spleen (**Fig. 1i**), were significantly lower compared to ND-fed mice. However, the lung weights (**Fig. S1a**) in both groups remained comparable.

**Figure 1:**
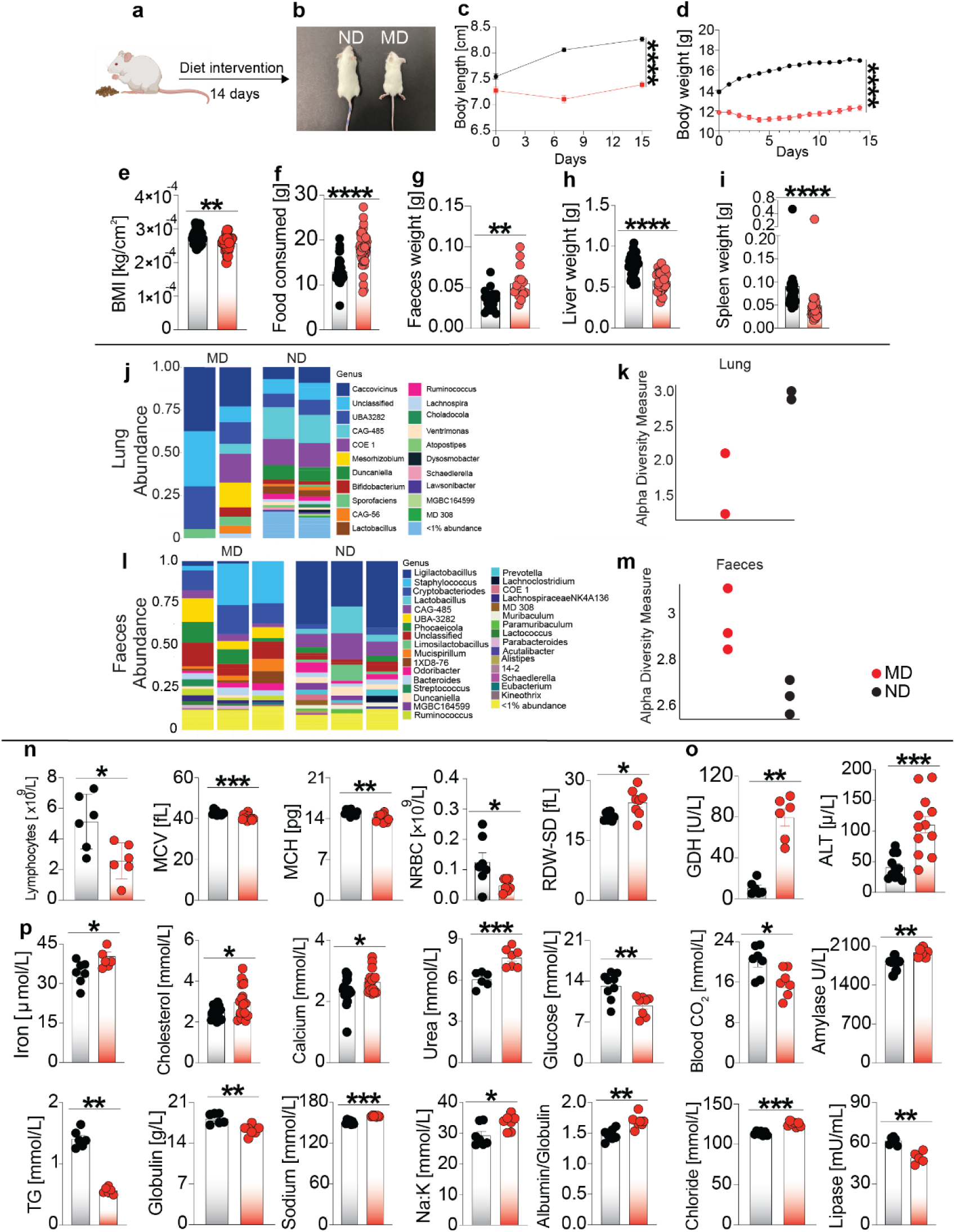
Characterisation of malnourished mice based on physical, haematological and biochemical parameters. **a)** Diet intervention scheme. **b)** Visual comparison between malnourished (right) and control (left) mice. **c)** Body length in cm. **d)** body weight in grams (black line: ND; red line: MD). **e)** Body Mass Index in kg/cm^2^; **f**) food consumption in gram; g) faeces pellet weight in gram; h) liver weight in gram; and **i**) spleen weight in gram at 14 days after diet intervention. **j, l)** Relative abundance of lung (**j**) and faeces (**l**) microbiota genera. **k, m)** Alpha diversity measure of lung (**k**) and faeces (**m**) microbiota. **n-p)** Key haematological (**n**) and biochemical (**o, p**) serum parameters at 15 days after diet intervention. Results are representative of five pooled independent experiments with 6 - 41 mice per group. Statistical significance was calculated using Mann-Whitney test (c-i, n-p); *p < 0.05, **p < 0.01, ***p < 0.001, ****p < 0.0001. MD, malnourished diet; ND, normal diet. Scheme in (a) generated with BioRender.

MD-fed mice exhibited lower lung microbial diversity (**Fig. 1j, k**) but increased gut microbial diversity than ND-fed mice, as demonstrated by the relative abundance and alpha diversity measure plots (**Fig. 1l, m**). Notably, microbial taxa including *Mucispirillum, Parabacteroides, Phocaeicola* and members of the *Lachnospiraceae* were uniquely observed in MD-fed mice, alongside microbiota shared with ND-fed mice. The lung microbial class abundance (**Fig. S1b, c**) and gut microbial class abundance (**Fig. S1d, e**) also differed between dietary groups.

Haematological analysis of blood revealed that, albeit not significant, the malnourished mice had nearly half the number of white blood cells (**Fig. S1f**) and lymphocytes (**Fig. 1n**, first plot). Additionally, malnourished mice showed slightly lower levels of haemoglobin **(Fig. S1g),** mean corpuscular volume (MCV) (**Fig. 1n**, second plot) and mean corpuscular haemoglobin (MCH) (**Fig. 1n**, third plot), and nucleated red blood cells (NRBC) (**Fig. 1n**, fourth plot). The mean corpuscular haemoglobin concentration (MCHC) (**Fig. S1h**), reticulocyte count (RET) (**Fig. S1i**) and platelets (PLT-F) (**Fig. S1j)** of malnourished and control mice remained similar.

Key liver enzyme activities also exhibited changes. Alanine aminotransferase (ALT) and glutamate dehydrogenase (GDH) levels were elevated in malnourished mice, indicating liver abnormalities (**Fig. 1o**, two plots). The malnourished mice also exhibited alterations in biochemical parameters (**Fig. 1p**). Elevated levels of iron, cholesterol, calcium and urea were observed (**Fig. 1p**, first four plots). However, malnourished mice had significantly low glucose and blood carbon dioxide levels than control mice (**Fig. 1p**, fifth and sixth plot) and comparable levels of creatinine and phosphorus (**Fig. S1k, l**) Additionally, malnourished mice had higher levels of amylase enzyme (**Fig. 1p**, seventh plot).

The triglyceride (TG) levels and globulin protein amount was found to be lower in malnourished mice compared to the normal diet group (**Fig. 1p**, eighth and ninth plots). The albumin and total protein content was similar in both groups (**Fig. S1m, n**). Electrolyte imbalances were also noted: sodium, sodium:potassium ratio and chloride levels were higher in malnourished mice (**Fig. 1p**, tenth, eleventh, thirteenth plot), while no significant change in potassium levels was noticed between the two groups (**Fig. S1o**). Malnourished mice also had a higher albumin-to-globulin ratio (**Fig. 1p**, twelfth plot). Lipase enzyme levels were lower in the MD-fed mice, suggesting pancreatic cell abnormalities in the malnourished mice (**Fig. 1p**, fourteenth plot).

Creatine kinase (CK) enzyme levels were slightly higher (insignificant) in MD-fed mice (**Fig. S1p**). Total bilirubin (TBIL) (**Fig. S1q**), total bile acids (TBA) (**Fig. S1r**) and direct bilirubin (DBIL) (**Fig. S1s**) also remained similar in malnourished and control mice. Other enzymes such as lactate dehydrogenase (LDH) (**Fig. S1t)**, alkaline phosphatase (ALP) **Fig. S1u)** and aspartate aminotransferase (AST) (**Fig. S1v)** remained comparable between the two groups. Collectively, those results indicated that MD-fed mice mimic key characteristics of human marasmus malnourishment.

### Malnourished mice are less susceptible to *Mtb* infection

To test the susceptibility of malnourished mice to *Mtb* infection and TB disease, mice were infected with a low infection dose (∼10 cfu) of *Mtb* H37Rv (**Fig. 2a**). Both groups inhaled a nearly equal number of tubercle bacilli illustrated by CFU measurements one day after infection (**Fig. 2b**). Interestingly, at 45 days after infection the bacterial burden in the lungs of malnourished mice was significantly lower compared to the ND group (**Fig. 2c**). This was reflected in the lung histopathological findings (**Fig. 2e, f**). There was no statistically significant difference in spleen CFU (**Fig. 2d**).

**Figure 2:**
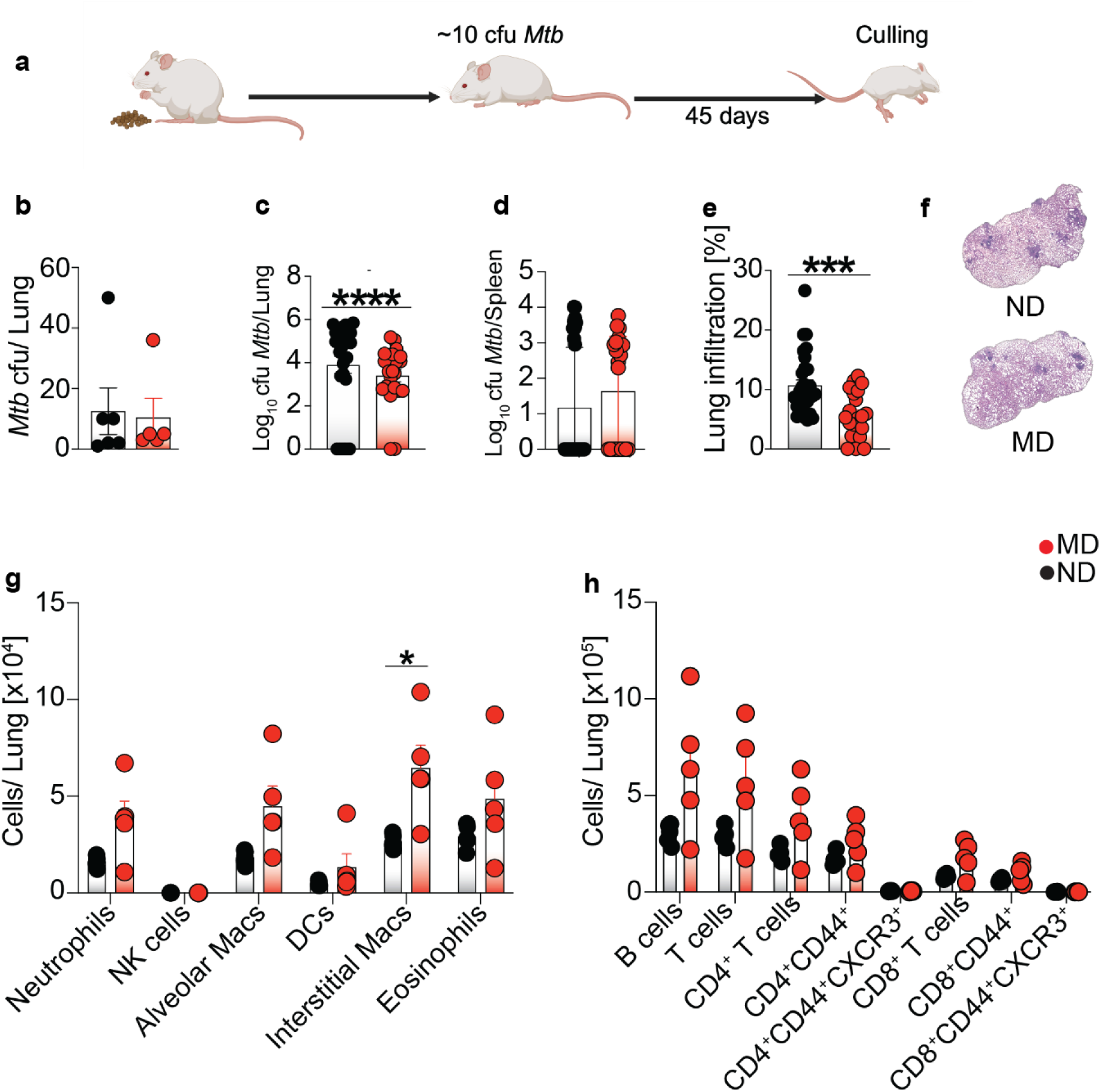
Susceptibility of malnourished mice to *Mtb* infection. **a)** Infection scheme. **b)** *Mtb* cfu in lungs one day after infection. **c-f)** Bacterial burden (log₁₀) in the lungs (**c)** and the spleen (**d**); Pulmonary tissue damage (**e)** and representative histological images (**f**). **g, h)** Flow cytometry results showing numbers of innate (**g**) and adaptive (**h)** immune cells in the lung. Results are representative of two pooled independent experiments except for FACS analysis. **b** (n = 6 mice per group), **c, d, e, f** (n = 22-30 mice per group), **g** and **h** (n = 5 mice per group). Statistical significance was calculated using Mann-Whitney test; *p < 0.05, **p < 0.01, ***p < 0.001, ****p < 0.0001. MD, malnourished diet; ND, normal diet. Scheme in (a) generated with BioRender.

While the reduced lung CFU burden was surprising, flow cytometry analysis of lung tissue revealed an overall elevated state of immune cell infiltration in MD-fed mice (**Fig. 2g, h**). Malnourished mice displayed a broad increase in CD4^+^ and CD8^+^ T cell subsets, including memory T cells (**Fig. 2h**). In addition, neutrophils, B cells, alveolar macrophages, dendritic cells (DCs), interstitial macrophages and eosinophils were increased in the lung of malnourished mice compared to the ND-fed group (**Fig. 2g**). Together, the results demonstrate that malnourished mice are less susceptible to TB disease and are characterised by an increased lung immune cell repertoire.

### BCG::ESAT-6-PE25SS maintained superior efficacy in malnourished mice

To assess the comparative efficacy of BCG and BCG::ESAT-6-PE25SS vaccine in malnourished mice, ND- and MD-fed mice were vaccinated intratrachealy (i.t.) with 10^5^ CFU and 60 days later were challenged with *Mtb* H37Rv (10-20 CFU) (**Fig. 3a**). The malnourished mice maintained significantly lower body weight throughout the study period, including after vaccination and after challenge (**Fig. 3b**), with significant differences observed at both the two-week (**Fig. 3c**) and 15-week (**Fig. 3d**) time points.

**Figure 3:**
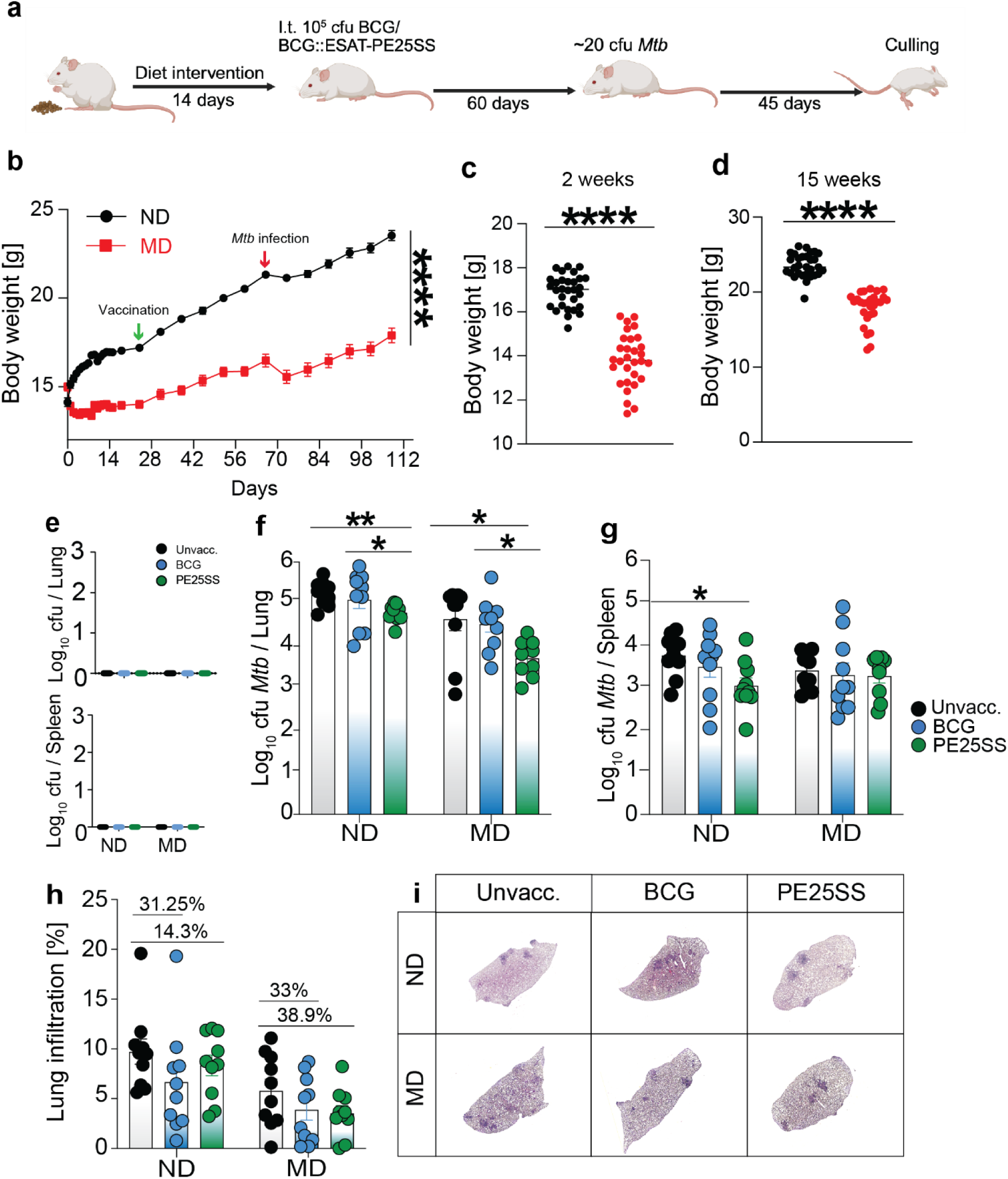
Evaluation of BCG::ESAT-6-PE25SS efficacy in malnourished mice. **a)** Schematic overview of the experimental design. **b)** Body weight over time. **c)** Body weight after 2 weeks on diet. **d)** Body weight after 15 weeks. **e)** Number of BCG and rBCG colonies in spleen and lung at 45 days after *Mtb* infecion. **f-h)** Bacterial burden (log_10_) in lung (**f**) and spleen (**g**), and lung infiltration (**h**), with percentage reduction relative to the control group indicated on each bar at 45 days after *Mtb* infection. **i)** Representative lung histopathology images. Results are representative of two pooled independent experiments. **b, c, d, e, f, g and h** (n = 9-10 mice per group) Statistical significance was calculated using Mann-Whitney test (**b, c, d**) ordinary one-way ANOVA (**e, f, g**); *p < 0.05, **p < 0.01, ***p < 0.001, ****p < 0.0001. MD, malnourished diet; ND, normal diet. Scheme in (a) generated with BioRender.

No residual vaccine bacilli were detected in the lung and the spleen post *Mtb* challenge (**Fig. 3e**). As expected, the unvaccinated mice in both nutritional groups exhibited the highest *Mtb* burden in the lungs (**Fig. 3f**). Notably, malnourished mice showed relatively lower lung CFU counts compared to the control group (**Fig. 3f**), consistent with earlier findings (**Fig. 2c**). Vaccination with BCG led to a modest reduction in lung bacterial burden of 0.21 and 0.11 log_10_ CFU compared to unvaccinated mice in both ND- and MD-fed mice, respectively. The most pronounced and significant reduction in lung bacterial load was observed in mice vaccinated with BCG::ESAT-6-PE25SS, compared to the unvaccinated or BCG-vaccinated control and malnourished mice (**Fig. 3f**).

In the spleen, the malnourished unvaccinated mice showed a lower *Mtb* burden compared to the control group (**Fig. 3g**). The BCG-vaccinated mice displayed a 0.27 log_10_ CFU reduction in ND mice and a 0.22 log_10_ CFU reduction in MD mice. Notably, ND-fed mice vaccinated with BCG::ESAT-6-PE25SS had the lowest spleen CFU counts with a 0.72 log_10_ CFU reduction, whereas the malnourished group vaccinated with BCG::ESAT-6-PE25SS showed CFU levels comparable to BCG vaccination with a 0.23 log_10_ CFU reduction.

Histopathology data (**Fig. 3h, i**) revealed notable differences between control and malnourished mice. Among the control mice, those vaccinated with BCG exhibited the least pulmonary damage, with a reduction of 31.25% compared to unvaccinated mice, followed by the group vaccinated with BCG::ESAT-6-PE25SS (14.29%). Consistent with the *Mtb* susceptibility data (**Fig. 2e, f**), the unvaccinated malnourished group had a lower percentage of lung damage compared to the unvaccinated ND group. Both vaccinated malnourished groups displayed similar levels of pulmonary damage, with BCG vaccination leading to 33% reduction in the lung infiltration and BCG::ESAT-6-PE25SS vaccination leading to 38.9% reduction (**Fig. 3h**). Collectively, those results indicate that BCG::ESAT-6-PE25SS vaccination largely maintains its superior efficacy in malnourished mice.

### Altered microbiota mice mimic human-like dysbiosis and microbial enrichment

To determine the role of an altered microbiome on *Mtb* susceptibility and vaccine-mediated protection, we first established ‘dirty’ mice and antibiotic-treated (ABX) cohorts (**Fig. 4a**) and characterised their microbial and cellular composition (**Fig. 4b-f**). Four to five months after co-housing of SPF mice with ‘dirty’ wild mice, and two weeks after antibiotic treatment, full-length 16S rRNA PacBio sequencing revealed a significant diversification of the lung and gut microbiota in SPF-Dirty mice and a near-absence of the detectable bacteria in ABX mice (**Fig. 4b, d**). SPF-Dirty mice displayed a similar Shannon diversity index to the two original Fight-4-Flight (F4F) and Berzerka (BZK) wild mice (**Fig. 4c**). FACS analysis of lung tissue revealed an increase in CD4^+^ and CD8^+^ T cell subsets in the lungs of SPF-Dirty mice (**Fig. 4g)** but similar cellular composition in ABX mice (**Fig. 4h**).

**Figure 4:**
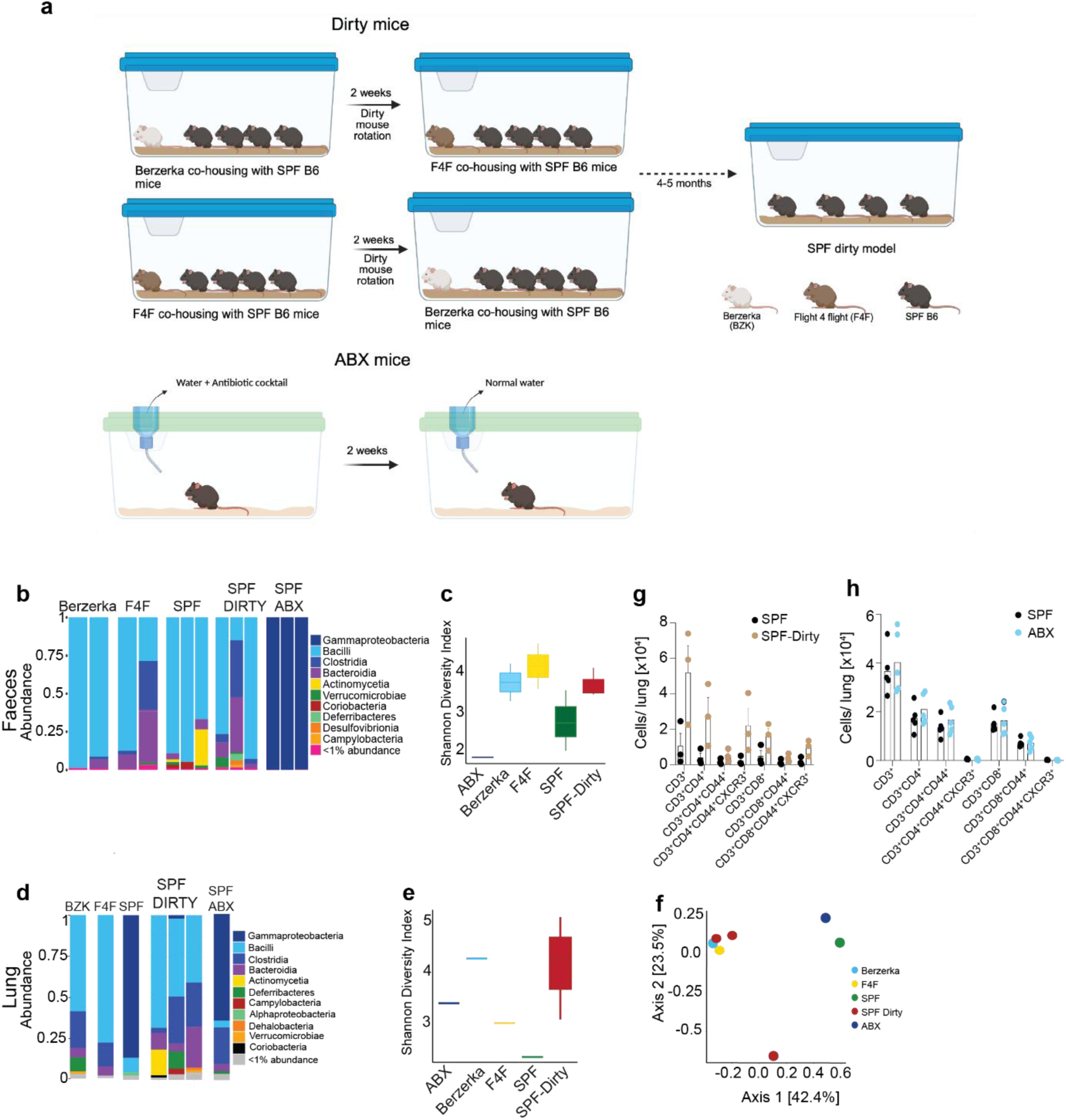
Characterisation of dirty and antibiotic-treated mice. **a)** Model development scheme. **b)** Relative abundance of faecal microbiota. **c)** Shannon index of faecal samples. **d)** Relative microbial abundance in lung tissue. **e)** Shannon index of lung microbiota. **f)** PCA plot of lung microbiota. **g, h)** FACS analysis of lung immune cells. Results are representative of one representative experiment. **b, c**, **f, g** and **h** (n = 2-3 mice per group) **d** and **e** (n = 5 mice per group). Statistical significance was calculated using Mann-Whitney test; *p < 0.05, **p < 0.01, ***p < 0.001, ****p < 0.0001. Scheme in (a) generated with BioRender.

Separate cohorts of SPF-Dirty and ABX mice were infected with about 20 cfu of *Mtb* to assess their susceptibility to TB (**Fig. 5a**). Mice from each model, along with their respective SPF control groups, inhaled a similar number of *Mtb* (**Fig. 5b, g**). At 45 days after *Mtb* challenge SPF-Dirty mice displayed reduced bacterial load in the lungs compared to the SPF group (**Fig. 5c**), while the bacterial counts in the spleen remained comparable between the two groups (**Fig. 5d**). Pulmonary damage in the ‘dirty’ mice was also reduced (16.11%) relative to the SPF group (**Fig. 5e, f**).

**Figure 5:**
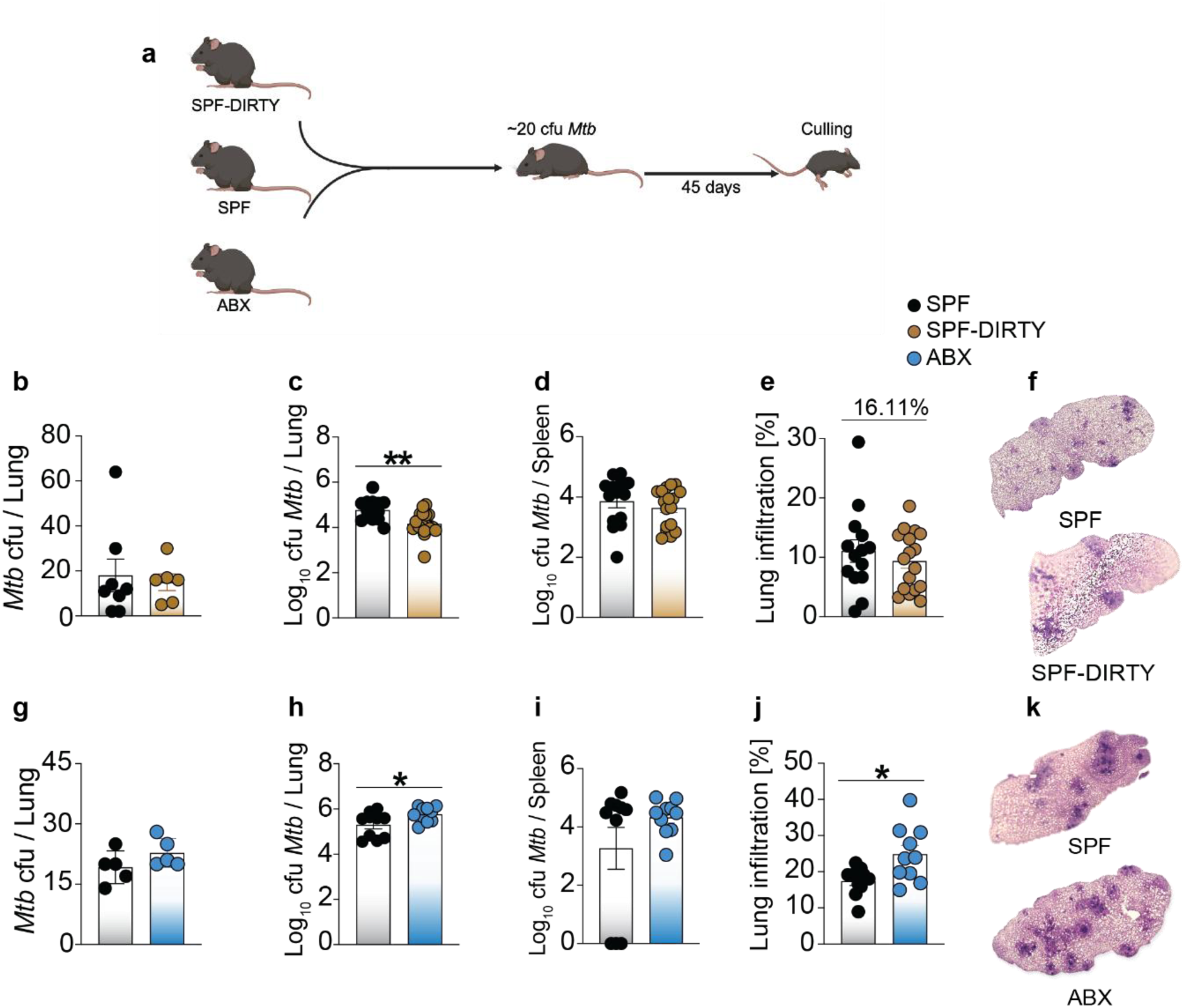
Susceptibility of dirty and antibiotic-treated mice to *Mtb* infection. **a)** Infection procedure scheme. *Mtb* cfu in lung at day 1 after infection of dirty mice (**b**) and ABX mice (**g**). **c-h; d-i)** Bacterial burden at 45 days after *Mtb* infection in the right lung lobe of dirty (**c**) and ABX mice (**h**); and in the spleen of dirty (**d**) and ABX mice (**i**). **e-j; f-k)** Percentage lung infiltration at 45 days after *Mtb* infection in dirty (**e**) and ABX mice (**j**) with a representative H&E image per group (**f, k**). Results are representative of two independent experiments. **a** (n=6 mice per group), **b, c, d** (n = 14-17 mice per group) **f** (n = 5 mice per group), **g, h, i** (n = 10 mice per group). Statistical significance was calculated using Mann-Whitney test; *p < 0.05, **p < 0.01, ***p < 0.001, ****p < 0.0001. Scheme in (a) generated with BioRender.

In contrast, ABX mice showed the opposite; with both lung CFU (**Fig. 5h**) and pulmonary cellular infiltration (**Fig. 5j, k**) being significantly increased compared to SPF controls. CFU levels in the spleen were also increased albeit not reaching statistical significance (**Fig. 5i**). Collectively, these results demonstrate that microbiome diversification is associated with increased resistance to TB, while microbial depletion leads to increased susceptibility.

### BCG::ESAT-6-PE25SS outperforms BCG efficacy in antibiotic-treated mice

To determine vaccine performance in ABX and SPF-Dirty mice, additional cohorts of mice were generated, vaccinated i.t. with BCG or BCG::ESAT-6-PE25SS and challenged with *Mtb* 60 days later (**Fig. 6a**). In the absence of vaccination, the antibiotic-treated group exhibited the greatest tubercle burden in both lungs (**Fig. 6b**) and spleen (**Fig. 6c**), followed by the unvaccinated SPF group. SPF-Dirty mice carried the lowest *Mtb* load, confirming results from our *Mtb* susceptibility experiments (**Fig. 5 c, e, f**). SPF mice immunized with BCG exhibited approximately 0.775 log_10_ fewer *Mtb* in the lung and 0.65 log_10_ fewer bacteria in the spleen, in line with expected efficacy of BCG in C57BL/6 mice^47^. BCG-mediated CFU reductions in ABX mice were slightly lower with a 0.03 log_10_ CFU reduction in the lung and a 1.04 log_10_ CFU reduction in the spleen. In SPF-Dirty mice there was no overall reduction seen in CFU with a large spread of CFU across individual mice. Vaccination with BCG::ESAT-6-PE25SS led to a significant reduction of 1.389 log_10_ relative to unvaccinated mice and a 0.614 log_10_ reduction relative to BCG vaccinated mice in the lung of SPF mice; and to a significant reduction of 0.6 log_10_ relative to unvaccinated mice and a 0.563 log_10_ reduction relative to BCG vaccinated mice in the lung of ABX mice (**Fig. 6b, c**). Similarly to BCG vaccination, there was no significant CFU reduction in SPF-Dirty mice vaccinated with BCG::ESAT-6-PE25SS vaccine.

**Figure 6:**
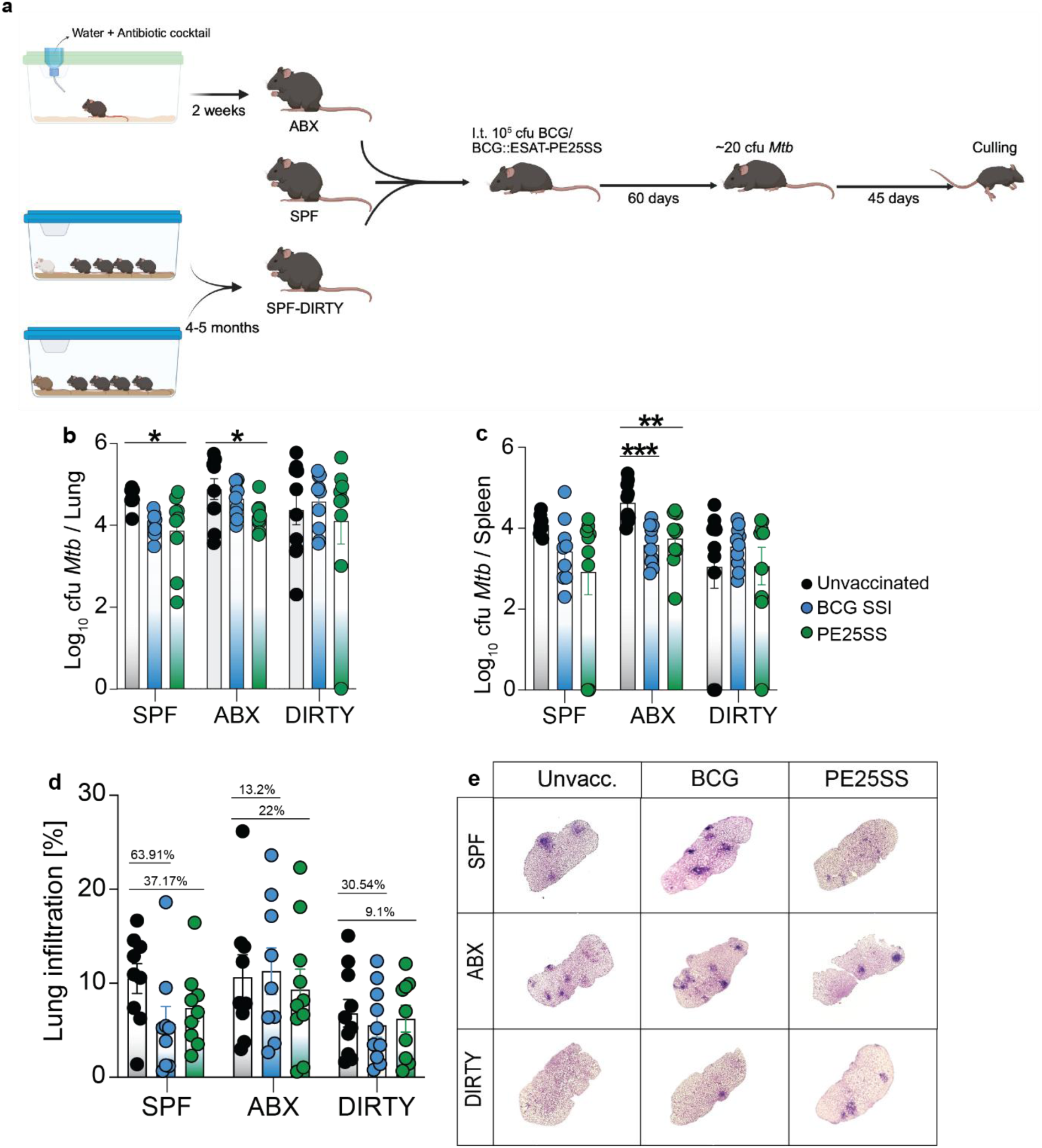
Efficacy of BCG::ESAT-6-PE25SS in dirty and antibiotic-treated mice. **a)** Overall scheme of experiment and induction of microbial alteration. **b-c)** *Mtb* cfu in lung (**b**) and spleen (**c**) across SPF, ABX and dirty mice at 45 days after *Mtb* infection. **d)** Percentage pulmonary infiltration in SPF, ABX and dirty mice at 45 days after *Mtb* infection, with percentage reduction relative to the control group indicated. **e)** Representative lung H&E images of SPF, ABX and dirty mice. Results are representative of two independent experiments. **b, c, d** (n = 9-10 mice per group) Statistical significance was calculated using ordinary one-way ANOVA; *p < 0.05, **p < 0.01, ***p < 0.001, ****p < 0.0001. Scheme in (a) generated with BioRender.

Pulmonary tissue damage (**Fig. 6d, e**) was most severe in the unvaccinated mice across all treatment groups. In SPF mice, BCG vaccination reduced lung infiltration by 63.91%, whereas BCG::ESAT-6-PE25SS vaccination resulted in a 37.17% reduction. In ABX mice, *Mtb*-induced tissue damage was increased by 13.2% following BCG vaccination and reduced by 22% following BCG::ESAT-6-PE25SS vaccination. Similarly, in SPF-Dirty mice, BCG vaccination reduced 30.54% pulmonary damage, whereas BCG::ESAT-6-PE25SS vaccination achieved a 9.1% reduction in lung infiltration (**Fig. 6d, e).**

Collectively, those results indicate that BCG::ESAT-6-PE25SS vaccination largely maintains its superior efficacy in antibiotic-treated mice and that overall vaccination efficacy in ‘dirty’ mice is very heterogenous, mimicking the variable efficacy of BCG vaccination observed in human populations.

### Baseline microbial profile dictates the host response to vaccination

Our results suggested that the microbiome composition in lung and gut may influence both susceptibility to TB disease as well as the host immune response to vaccination. To determine how microbial ecology influences TB susceptibility and vaccine responses, we profiled gut and lung microbiota by 16S rRNA sequencing in SPF, ABX, and SPF-Dirty mice at baseline (B), post-vaccination (V), and post-*Mtb* infection (I) (**Fig. 7**). As expected, baseline gut microbial diversity was highest in SPF-Dirty mice, intermediate in SPF, and profoundly reduced in ABX animals, which exhibited a collapsed microbiota dominated by residual taxa (**Fig. 7a, b, c).** The residual bacterial genera in ABX mice post-treatment were predominantly resistant to the antibiotics used **(Fig. 7d).**

**Figure 7:**
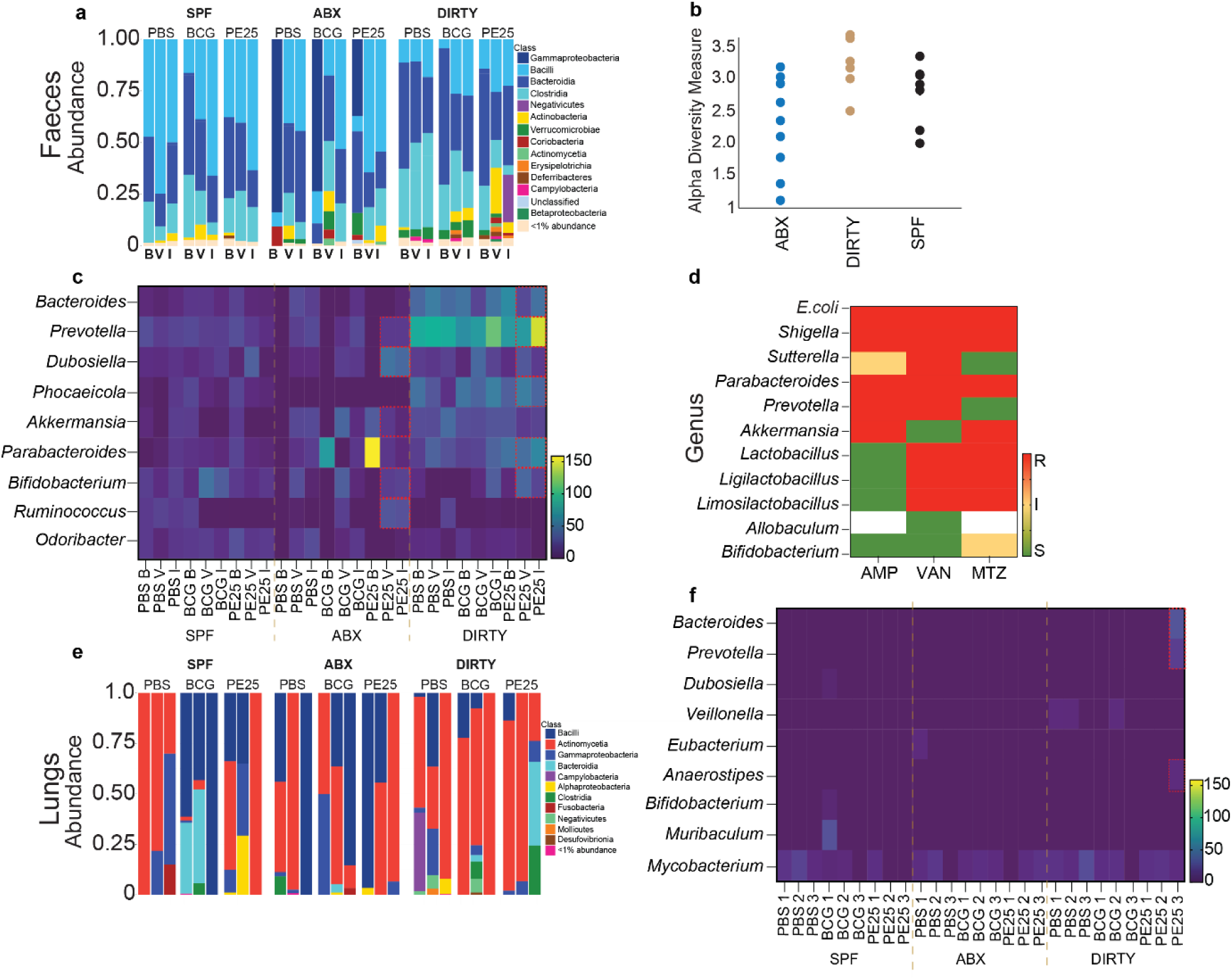
Microbial diversity following vaccination and challenge across SPF, ABX and dirty mice. **a)** Relative abundance of bacterial classes in faeces. **b)** Alpha diversity measure across bacterial genera in faeces. **c**) Heatmap of relative abundance of selective genera able that produce short-chain fatty acids (SCFA) in gut and their abundance before and after vaccination, and after *Mtb* challenge. **d**) Heatmap of genera present post-antibiotic treatment. **e**) Relative abundance of bacterial classes in lungs. **f**) Heatmap of relative abundance of SCFA-producing genera in lung and their abundance before and after vaccination, and after *Mtb* challenge. Results are representative of one experiment. **a, b, c, d, e, f** (n = 5 mice per group). B, baseline; V, 60 days after vaccination; I, 45 days after *Mtb* infection. R, resistant; I: intermediate and S: susceptible.

Vaccination induced a pronounced and vaccine-dependent microbial shift (**Fig. 7c).** Notably, BCG::ESAT-6-PE25SS vaccination consistently drove the strongest and most distinct microbial remodelling across models, characterised by expansion of *Prevotella*, *Dubosiella*, *Bifidobacterium and Ruminococcus* in ABX and *Bacteroides, Prevotella, Dubosiella, Phocaeicola, Parabacteroides* and *Bifidobacterium* in ‘dirty’ mice (**Fig. 7a, c and S2a)**. In contrast, BCG vaccination partially increased the proportion of *Dubosiella* and *Akkermansia* populations in ABX mice and more modest changes in SPF-Dirty animals (**Fig. 7a, c and S2a)**.

Following *Mtb* infection, microbial trajectories diverged further by vaccine type (**Fig. 7a, c**). In ABX mice, unvaccinated animals exhibited a collapse of *Bifidobacterium*, whereas BCG vaccination resulted in loss of key commensals such as *Bifidobacterium* and *Akkermansia*. BCG::ESAT-6-PE25SS vaccination on the other hand preserved and/or expanded multiple commensals, including *Bacteroides, Prevotella, Dubosiella, Akkermansia, Parabacteroides, Bifidobacterium*, *Ruminococcus* and *Odoribacter*, indicating enhanced microbiome resilience in this group during *Mtb* infection (**Fig. 7a, c and S2a)**.

Lung microbiome profiling revealed lower diversity but mirrored the trend observed in gut microbiome (**Fig. 7e, f**). Across models, BCG::ESAT-6-PE25SS sustained a distinct lung microbial signature, including enrichment of *Bacteroides, Prevotella and Anaerostipes,* in ‘dirty mice*’,* differentiating it from BCG- or unvaccinated groups (**Fig. 7e, f and S2b)**. **Fig. S3 and S4** present bidirectional plots illustrating the mean percentage abundance of bacterial taxa across the different groups. Overall, it appeared that microbiome complexity may influence ecological resilience to *Mtb* infection, while BCG::ESAT-6-PE25SS vaccination consistently induced the most pronounced and stable microbial reconfiguration across both gut and lung compartments, particularly in ABX and SPF-Dirty groups.

Correlation analyses between lung CFU levels and gut microbial abundance (**Fig. 8 and Fig. S5**) further revealed distinct associations between gut microbial genera and pulmonary *Mtb* burden across vaccination status and microbiome composition (correlation analyses with lung microbial diversity could not be performed because of the low sequencing depth and limited read recovery in lung samples). The BCG::ESAT-6-PE25SS vaccination group demonstrated stronger microbiota-CFU directional trends compared with BCG, suggesting enhanced microbiome-dependent immune modulation. Particularly, the abundance of genera such as *Bacteroides, Prevotella, Phocaeicola, Akkermansia and Parabacteroides* in BCG::ESAT-6-PE25SS vaccination group demonstrated the strongest associations, exhibiting the highest R^2^ values, compared with BCG across the ABX, dirty and SPF groups (**Fig. 8**). ‘Dirty’ mice exhibited the most pronounced bacterial-CFU relationships, supporting a role for microbiome complexity in shaping anti-mycobacterial immunity.

**Figure 8:**
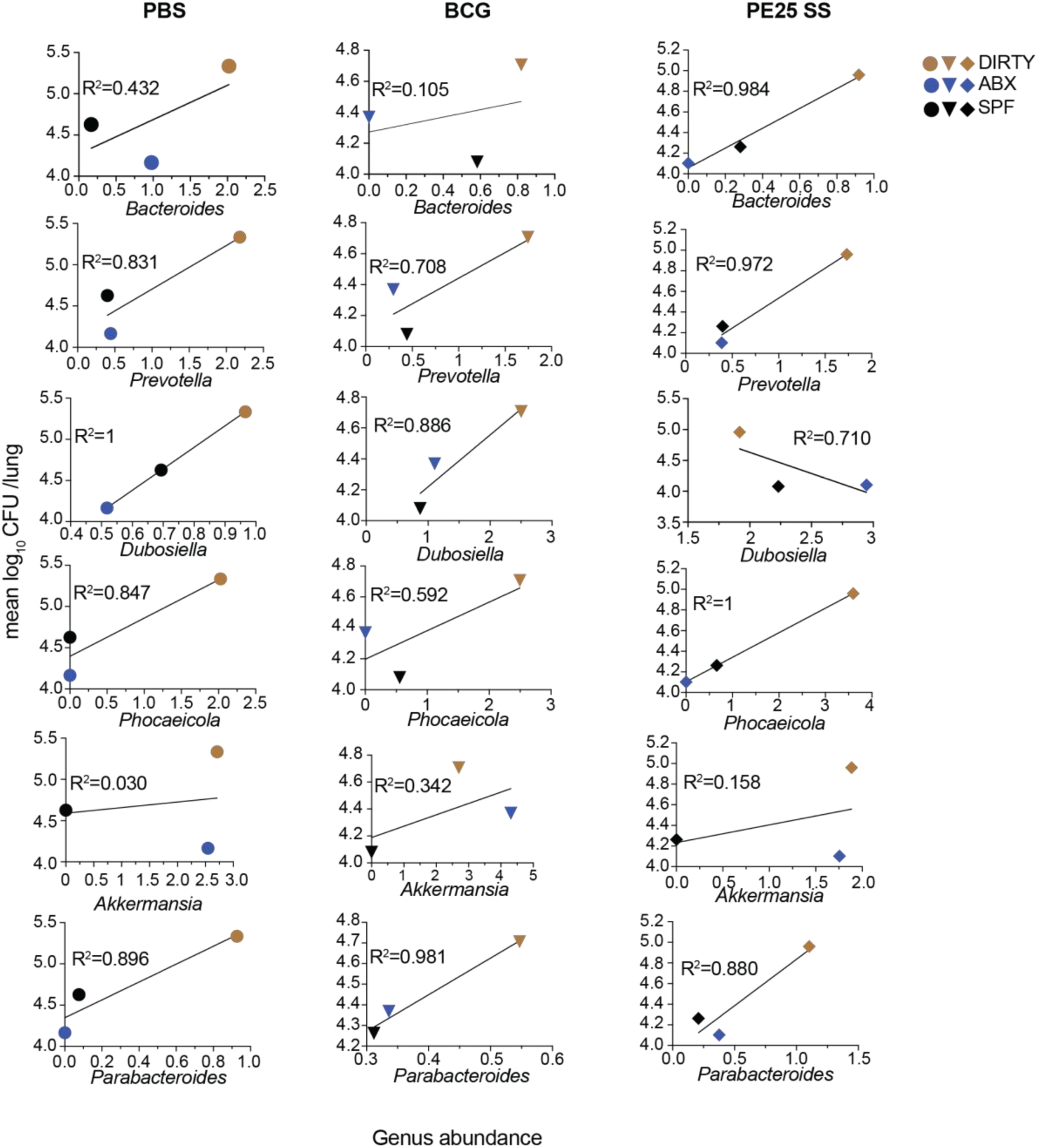
Correlation between lung *Mtb* burden and bacterial abundance. Mean *Mtb* CFU versus bacterial abundance plots, including R^2^ values, for selected SCFA-producing bacterial genera (*Bacteroides, Prevotella, Dubosiella, Veillonella, Akkermansia and Parabacteroides)* in unvaccinated (left), BCG-vaccinated (middle) and BCG::ESAT-6-PE25SS-vaccinated (right) groups.

## Discussion

Here we investigated how dietary composition and gut microbiota influence host immunity, susceptibility to *Mtb*, and the efficacy of a recombinant BCG vaccine candidate. Our findings demonstrate that nutritional status and microbiome composition substantially reshape immune responses and *Mtb* control.

Unexpectedly, marasmus-like malnourished mice exhibited reduced pulmonary *Mtb* burden and lower lung pathology compared to control animals. At first glance, this contrasts with the established association between malnutrition and increased infection susceptibility in humans^48–50^. Our results suggest that dietary composition and metabolic adaptation can reprogram host immunity in ways that restrict pathogen growth. Alongside increased T cell numbers, one key observation was the expansion of innate immune populations, particularly interstitial macrophages, which are associated with glycolytic metabolism and enhanced intracellular restriction of *Mtb*^51^. Additionally, dietary composition, which was high in carbohydrates but low in fat and protein, may have altered host lipid metabolism, reducing availability of triglycerides and intracellular cholesterol-rich niches required for *Mtb* survival. Since *Mtb* relies on host lipids^52^ and foamy macrophages during early infection^52, 53^, reduced lipid availability may have imposed metabolic stress on *Mtb*^54^. Together, these changes suggest that malnutrition in this model does not simply suppress immunity but induces a distinct metabolic-immune state that can restrict *Mtb* growth. It is well established that calorie restriction in humans and animals, for example via intermittent fasting, leads to increased energy metabolism and activated immunity^55, 56^. The malnourished diet used in this study partially mimics the vegetarian/vegan and Mediterranean diet^57, 58^. It is tempting to speculate that in such environments, distinct vaccine candidates may help elicit improved immune responses against TB.

Beyond diet, microbiome composition emerged as a major determinant of TB susceptibility. Antibiotic-treated mice^38^ with disrupted microbiota exhibited increased bacterial burden and more severe pathology, whereas ‘dirty’ mice with environmentally enriched microbiota showed enhanced resistance to infection. Dirty mice displayed broader T cell repertoires and immune phenotypes resembling more mature human immune systems, supporting the idea that microbial exposure enhances immune readiness^40, 41^. Importantly, vaccination with BCG or rBCG reduced disease severity even in dysbiotic conditions, although efficacy varied with microbiome context. While the malnourished mice were not included in microbiota-based vaccine efficacy comparisons, dietary intervention alone significantly altered microbial diversity and immune priming. This suggests that nutritional status and microbiome composition independently and interactively shape host responses to *Mtb* and vaccination.

Across models, BCG::ESAT-6-PE25SS^30^ demonstrated superior protection compared to conventional BCG, with significantly reduced pulmonary *Mtb* burden and improved histopathological outcomes. The enhanced efficacy was maintained under malnourished, and antibiotic-treated conditions, suggesting robustness across diverse immune states. While a detailed immunological analysis was beyond the scope of this study, the secretion of ESAT-6 likely enhanced immune recognition via cytosolic sensing pathways such as cGAS, AIM2, and NOD-like receptors, amplifying both innate and adaptive responses^59^.

Interestingly, microbiome profiling revealed that vaccination with rBCG was associated with enrichment of SCFA-producing microbial taxa, including *Bacteroides, Prevotella, Phocaeicola*, *Parabacteroides*, *Dubosiella, Akkermansia, Bifidobacterium and Ruminococcus.* SCFAs are known to regulate immune function through G-protein–coupled receptors and epigenetic modulation, promoting macrophage activation and balanced inflammatory responses^60^. While conventional BCG vaccination was associated with butyrate-producing microbial shifts, rBCG favoured propionate-associated communities (*Parabacteroides, Prevotella, Veillonella and Phocaeicola*) and cross-feeders (*Dubosiella*) potentially contributing to differential immune programming^61^.

These microbiome-driven effects suggest a functional gut-lung axis influencing vaccine efficacy. It is known that SCFA signalling through receptors such as GPR41 and GPR43 modulates macrophage metabolic states and enhances pathogen control while limiting immunopathology^59, 62^. BCG::ESAT-6-PE25SS appeared to induce greater microbiome modulation than BCG, suggesting a stronger microbiota-dependent immunomodulatory effect. Our results imply that the expanded antigenic repertoire of a live recombinant vaccine may interact differently with the host microbiota, thereby reshaping the microbial niche and subsequently impacting the host immune system to achieve improved control of *Mtb* infection. The magnitude of such vaccine-microbiota interactions and their downstream immunological effects is likely less pronounced in subunit or multiepitope peptide candidates due to their relative limited antigenic complexity. However, this hypothesis requires experimental validation.

Importantly, our findings highlight limitations of conventional SPF mouse models used in vaccine development. These animals possess relatively limited microbiota complexity and immature immune systems, resembling human infancy rather than adult immune conditions^41^. In contrast, models with environmental microbial exposure or dysbiosis more closely reflect human heterogeneity, particularly in low- and middle-income countries where TB burden is highest. All three models used here: i) malnutrition, ii) antibiotic-induced dysbiosis, and iii) environmental microbiome exposure mirror clinically important human conditions. Marasmus reflects chronic energy deficiency and immune suppression; antibiotic-treated models mimic clinical microbiome disruption; and ‘dirty’ mice represent adult-like microbial complexity.

Given the repeated failure of advanced TB vaccine candidates in late-stage clinical trials, incorporating such physiologically relevant TB animal models that better reflect real-world conditions in preclinical testing would provide a crucial insight for the evaluation of the vaccine candidates under development and may improve translational relevance.

Several limitations of the study should be acknowledged. The functional phenotypes of macrophage subsets were not directly characterised, and future studies employing metabolic or single-cell transcriptomic approaches and cytokine analysis will be necessary to define the precise cellular mechanisms underlying *Mtb* restriction in these models. In addition, although associations between microbiota composition and vaccine efficacy were observed, causal relationships remain to be established. Furthermore, the published diet^63^ used to develop the malnourished mice contained high levels of micronutrients such as iron, zinc, copper, and manganese. These micronutrients play pivotal roles in controlling *Mtb*^64^. A more comprehensive evaluation of their elevated levels would provide deeper insights into the results obtained in the malnourished mice.

In conclusion, our study shows that diet, microbiome composition, and host metabolism jointly regulate immune responses to *Mtb* and significantly influence vaccine efficacy. The recombinant BCG::ESAT-6-PE25SS vaccine consistently outperformed conventional BCG across diverse host conditions, suggesting improved robustness in real-world settings. These findings emphasize the need to integrate nutritional and microbial context into TB vaccine development and highlight the gut-lung axis as a key determinant of protective immunity. Future studies should focus on mechanistic dissection using microbiota transfer, metabolite supplementation, and single-cell immune profiling to establish causal pathways underlying these observations.

## Methods

### Ethics statement

This study was conducted in accordance with the National Health and Medical Research Council (NHMRC) animal care guidelines with all procedures approved by the animal ethics committee (A2855 and A2865) of James Cook University, Australia.

### Mice

BALB/c and C57BL/6J were obtained from the Animal Resources Centre (ARC) or the Australian Bio Resources (ABR). Upon arrival, mice were housed in a specific pathogen-free (SPF) animal facility within the Australian Institute of Tropical Health and Medicine (AITHM), James Cook University, Townsville, Australia. The original ‘dirty’ wild mice, named Berzerka (BZK) and Fight 4 Flight (F4F) of unknown age were obtained from two different reptile food vendors based in Townsville, Australia.

#### Malnourished model

Female BALB/c mice of 4-5 weeks of age were randomly assigned to two groups: normal diet (ND) and malnourished diet (MD). The ND group received a standard diet ad libitum (SF08-020, Specialty Feeds, Western Australia), while the MD group was fed with a customized malnourished diet (SF22-108, modified NIH-31; energy restricted protein 8% and fat 2.2%, Specialty Feeds, Western Australia) as reported previously^63^. Both groups received 30 g of feed per cage containing five mice daily. The malnourished diet consisted of high carbohydrate, high fibre, low fat, and low protein content. For model characterization, both groups were maintained on their respective diets for 14-16 days. Mice were then euthanized, and blood samples were collected for hematology and biochemical analyses. Organs including lungs, liver, and spleen were also collected and weighed post-diet intervention.

#### Antibiotic-treated model

Female 4–5-week-old C57BL/6J mice were used for the antibiotic treatment. A cocktail of ampicillin (AMP;1 g/L), metronidazole (MTZ; 1 g/L), and vancomycin (VAN; 0.5 g/L) was added to sterile reverse osmosis (RO) water supplemented with 3% sucrose to mitigate reduced water intake due to the bitter taste of the drugs^38, 65^. Mice received the medicated water for two weeks, after which they were euthanized for verification of microbiome composition.

#### Dirty mouse model

The ‘dirty’ mice were treated with ivermectin to prevent pinworm contamination prior to use in experiments. Female 5-week-old C57BL/6J SPF mice were co-housed with either Berzerka or Fight 4 Flight at a ratio of 4:1. To promote microbial diversity, ‘dirty mice’ were rotated between cages every two weeks. Co-housing continued for 3-5 months. All mice, including the original wild ‘dirty’ mice, were subsequently euthanized for analysis to confirm microbial diversity.

### Bacterial strains

BCG SSI, recombinant BCG::ESAT-6-PE25SS, and *Mtb* H37Rv were cultured in Middlebrook 7H9 broth (BD Biosciences). The medium was supplemented with 0.2% glycerol, 0.05% Tween-80 (Sigma-Aldrich), 10% albumin dextrose catalase (ADC) enrichment (BD Biosciences), and appropriate antibiotics as required. Cultures were harvested at mid-logarithmic phase (OD_600_-0.6–0.8), washed in sterile phosphate-buffered saline (PBS), aliquoted into 1 mL cryovials, and stored in PBS containing 15% glycerol at -80°C until required for vaccination or infection. The number of colony-forming units (CFUs) in the inoculum was determined prior to use.

### Vaccination and infection

For vaccination and infection, cryopreserved bacterial stocks were thawed, centrifuged, and the pellets resuspended in sterile PBS to achieve the appropriate dose. All mice were immunized under BSL-2 conditions via intratracheal (i.t.) administration of 5 × 10^5^ CFUs per mouse. Mice were anesthetized using a portable isoflurane system (3-5% for induction, 3% for maintenance). The tongue was gently retracted using sterile blunt forceps, and 50 µL of inoculum was administered into the oropharynx using a 200 µL pipette. The nostrils were briefly occluded to ensure delivery of the inoculum into the lungs.

Sixty days post-vaccination (p.v.), mice were challenged with a low aerosol dose of *Mtb* H37Rv (10–20 CFUs) using a Glas-Col inhalation exposure system within a BSL-3 suite. The initial infectious dose was verified by homogenizing lung tissue from five mice at one day post-infection (p.i.), followed by plating on 10% OADC-enriched 7H11 agar.

### Sample collection

Forty-five days post-infection, mice were euthanized by cervical dislocation. Blood was collected via cardiac puncture and transferred to Z-gel tubes (Sarstedt) for centrifugation to obtain serum. Serum was filtered through a 0.2 µm SpinX column (Sigma) and stored at -80°C. Organs including lungs and spleens were aseptically harvested for CFU enumeration and histological analysis.

### CFU enumeration

Lung (right lobe) and spleen were homogenized in gentleMACS tubes containing 1 mL sterile PBS with 0.05% Tween-80 using a tissue homogenizer. Serial dilutions of the homogenates were prepared in a 45-well polystyrene plate containing 900 µL of PBS/0.05% Tween-80 per well. Aliquots (100 µL) were plated onto 10% oleic albumin dextrose catalase (OADC)-enriched 7H11 agar supplemented with 10 mg/mL cycloheximide and 25 µg/mL ampicillin for *Mtb*, or 50 µg/mL kanamycin for recombinant BCG. Sterility testing of the BCG strain was performed on the plates without antibiotics. Plates were sealed and incubated aerobically at 37 °C for 3-4 weeks. Colonies were counted, and total CFUs per organ were calculated based on dilution factors.

### Lung histology

Left lung lobes were collected for histopathological evaluation. Tissues were fixed overnight in 4% paraformaldehyde, transferred into cassettes, and stored in 70% ethanol until processing. Fixed tissues were embedded in paraffin, and 4 µm sections were prepared using a microtome. Sections were mounted on labelled glass slides and stained with haematoxylin and eosin (H&E). Quantification of lung pathology was performed using ImageJ, National Institutes of Health (NIH) software by calculating the percentage of tissue area affected, based on total lung area and regions showing dense cellular infiltration, as previously described^11^.

### Haematology and biochemical analysis

Blood was collected via terminal cardiac puncture while the mice were under anaesthesia. The liver, lungs, and spleen were also excised for weight measurement. A 1 mL syringe fitted with a 25G needle was used for blood collection, and the syringe was pre-lubricated with freshly prepared sterile EDTA solution for haematology. Blood was collected in EDTA tubes, while serum tubes were used for biochemical testing. The serum tubes were centrifuged to obtain serum. All collected samples were stored on ice and immediately transported to the Veterinary Diagnostic Laboratory at James Cook University, Townsville.

### Flow cytometry

The lung tissue was homogenized in gentleMACS tubes containing RPMI, followed by brief centrifugation. Subsequently, 2 mL of collagenase solution was added, and the mixture was incubated at 37°C for 30 minutes for enzymatic digestion. After incubation, the solution was passed through a cell strainer using a syringe plunger into a 50 mL tube. The cell suspension was centrifuged, and the pellet resuspended in red blood cell lysis buffer. Following lysis, the tube was topped up with FACS buffer and centrifuged again. Finally, the cell pellet was resuspended in FACS buffer for antibody staining. The following antibodies from BD Biosciences were used CXCR3 (BV788), CD19 (PE-CF594), F4-80 (BV650), CD44 (BV421), CD8 (BV510), CD335 (NKp46; BV711), Ly6G (BUV395), CD3 (Alexa700), CD206 (APCAF647), CD11c (PE-Cy7), fixable viability dye (APC-Cy7), CD4 (PerCP Cy5.5), Siglec F (PE), CD16/CD32 (Fc-Block).

### 16S rRNA sequencing

Mice with depleted (antibiotic-treated) or enhanced (‘dirty’) microbiota were euthanized for collection of lungs (n = 2-3 per group) and faecal samples (n = >5 per group) to verify successful model development. All samples were transferred into cryotubes and immediately stored at -80°C until further processing. The faecal samples from vaccination experiment were processed and stored in same way.

Lung tissues from vaccination experiments were processed for genomic DNA extraction using the PureLink™ Microbiome DNA Purification Kit (Invitrogen™). Initial tissue dissociation was performed in a BSL-3 laboratory using gentleMACS tubes and a tissue dissociator (Miltenyi Biotec) prior to genomic DNA extraction and sequencing submission. Faecal samples were subsequently submitted to the Australian Genome Research Facility (AGRF) for full length 16S PacBio sequencing.

The 16S rRNA PacBio sequencing targeted all variable regions (V1-V9) in bacterial 16S rRNA gene using a primer set F27 (GCATC/barcode/AGRGTTYGATYMTGGCTCAG) and R1492 (GCATC/barcode/RGYTACCTTGTTACGACTT). The data was quality filtered, primer trimmed and denoised to generate high-quality amplicon sequence variants (ASVs) using QIIME2^66^ and DADA^67^. Taxonomic classification was performed using the naïve Bayesian classifier implemented in DADA2 against three reference databases queried sequentially. Species-level assignments were first performed using the Genome Taxonomy Database (GTDB r207)^68^, followed by the SILVA rRNA database (v138)^69^ and finally the NCBI RefSeq 16S rRNA database supplemented with the Ribosomal Database Project (RDP)^70^ when no species-level match was identified. This hierarchical approach improved the taxonomic classification of low-abundance ASVs. The ASV tables and taxonomic annotations were subsequently exported in BIOM format for downstream analysis.

For the dirty mouse dataset (n=2-3), 384,861 PacBio HiFi full-length 16S rRNA reads were generated. Following quality filtering and primer removal, 378,990 reads were retained, of which 245,039 reads were used to generate 1,369 ASVs. For faecal samples (n= 27), 2.35 million PacBio HiFi reads were generated, with 1.79 million reads retained after quality and primer filtering. A total of 804,121 reads were subsequently used to generate 2,814 ASVs. For lung samples (n=27), 158,931 PacBio HiFi reads were generated, with 82,101 reads retained following quality and primer filtering. Of these, 72,363 reads were used to generate 279ASVs. The resulting BIOM files were analysed using the ConsensusMetaDA package (v1.0)^71^ in R version 4.5.1^72^. ConsensusMetaDA integrates several R packages, including phyloseq (v1.5.1)^73^, dplyr (v1.1.4)^74^ and ggplot2 (v4.0)^75^, for microbiome data analysis and visualization. Alpha-diversity analysis, ordination plots, and taxonomic abundance visualizations were generated using ConsensusMetaDA. Correlation analyses were performed using the mean genus-level abundance and corresponding lung CFU data.

### Statistics

Statistical analyses were performed, and graphs were generated using Prism version 11.0.2 (GraphPad). Two and multiple parametric group analyses were carried out using Mann-Whitney and one-way analysis of variance (ANOVA), respectively. p < 0.05 was considered significant, unless otherwise stated, *p < 0.05, **p < 0.01, ***p < 0.001, ****p < 0.0001.

## Acknowledgements

We thank Dr Helma Antony, Leanne Taylor and Dr Socorro Miranda-Hernandez for their support in the BSL2 and BSL3 laboratory, Olivia Johnson for her help in animal facility, and Erin Roberts and Roselfina Charol for their assistance in the histology lab.

## Author contributions

MP and AK conceptualized the study. MP, SR, KR, MK, HN, HS and AK performed experiments and analysed data. RR developed the dirty mouse model. JW assisted in developing ABX model with antibiotic treatment schedules and antibiotic selection. MP wrote the first manuscript draft. AK and MP edited and finalised the manuscript. SS, JW, CR and MAF provided editorial and intellectual input. All authors contributed to manuscript revision, read, and approved the submitted version.

## Data Availability

Requests for further information and resources should be directed to and will be fulfilled by the lead contact, Andreas Kupz. This study did not generate new, unique reagents. All data reported in this paper will be shared by the lead contact upon request.

## Code availability

This paper does not report original codes. Any additional information required to reanalyze the data reported in this paper is available in the main text and supplemental information or from the lead contact upon request.

## REFERENCES

1. WHO. Global Tuberculosis Report. (2025).

2. Creswell, J. et al. Tuberculosis and noncommunicable diseases: neglected links and missed opportunities. Eur Respir J 37, 1269–1282 (2011).

3. Kaluvu, L. et al. Multimorbidity of communicable and non-communicable diseases in low- and middle-income countries: A systematic review. J Multimorb Comorb 12, 26335565221112593 (2022).

4. Borges, A.H. et al. Immunogenicity, safety, and efficacy of the vaccine H56:IC31 in reducing the rate of tuberculosis disease recurrence in HIV-negative adults successfully treated for drug-susceptible pulmonary tuberculosis: a double-blind, randomised, placebo-controlled, phase 2b trial. Lancet Infect Dis 25, 751–763 (2025).

5. Udoakang, A.J. et al. The COVID-19, tuberculosis and HIV/AIDS: Ménage à Trois. Front Immunol 14, 1104828 (2023).

6. Lu, P. et al. Evaluation of ESAT6-CFP10 Skin Test for Mycobacterium tuberculosis Infection among Persons Living with HIV in China. Journal of Clinical Microbiology 61 (2023).

7. Girishbhai Patel, D., et al. Nutritional status in patients with tuberculosis and diabetes mellitus: A comparative observational study. Journal of Clinical Tuberculosis and Other Mycobacterial Diseases 35 (2024).

8. Boadu, A.A., Yeboah-Manu, M., Osei-Wusu, S. & Yeboah-Manu, D. Tuberculosis and diabetes mellitus: The complexity of the comorbid interactions. Int J Infect Dis 146, 107140 (2024).

9. Wang, Q., Ma, A., Schouten, E.G. & Kok, F.J. A double burden of tuberculosis and diabetes mellitus and the possible role of vitamin D deficiency. Clin Nutr 40, 350–357 (2021).

10. Sathkumara, H.D. et al. A murine model of tuberculosis/type 2 diabetes comorbidity for investigating the microbiome, metabolome and associated immune parameters. Animal Model Exp Med 4, 181–188 (2021).

11. Sathkumara, H.D. et al. Mucosal delivery of ESX-1-expressing BCG strains provides superior immunity against tuberculosis in murine type 2 diabetes. Proc Natl Acad Sci U S A 117, 20848–20859 (2020).

12. Alim, M.A. et al. Increased susceptibility to Mycobacterium tuberculosis infection in a diet-induced murine model of type 2 diabetes. Microbes Infect 22, 303–311 (2020).

13. Castro, D.B., Pinto, R.C., Albuquerque, B.C., Sadahiro, M. & Braga, J.U. The Socioeconomic Factors and the Indigenous Component of Tuberculosis in Amazonas. PLoS One 11, e0158574 (2016).

14. Wang, Q. et al. Spatial distribution of tuberculosis and its socioeconomic influencing factors in mainland China 2013-2016. Trop Med Int Health 24, 1104–1113 (2019).

15. Rafiq, M., Saqib, S.E. & Atiq, M. Health-Related Quality of Life of Tuberculosis Patients and the Role of Socioeconomic Factors: A Mixed-Method Study. Am J Trop Med Hyg 106, 80–87 (2021).

16. Dias, S., Castro, S., Ribeiro, A.I., Krainski, E.T. & Duarte, R. Geographic patterns and hotspots of pediatric tuberculosis: the role of socioeconomic determinants. J Bras Pneumol 49, e20230004 (2023).

17. Taal, A.T. et al. The geographical distribution and socioeconomic risk factors of COVID-19, tuberculosis and leprosy in Fortaleza, Brazil. BMC Infect Dis 23, 662 (2023).

18. Jesus, G.S. et al. Effects of conditional cash transfers on tuberculosis incidence and mortality according to race, ethnicity and socioeconomic factors in the 100 Million Brazilian Cohort. Nat Med 31, 653–662 (2025).

19. Morales, F., Montserrat-de la Paz, S., Leon, M.J. & Rivero-Pino, F. Effects of Malnutrition on the Immune System and Infection and the Role of Nutritional Strategies Regarding Improvements in Children’s Health Status: A Literature Review. Nutrients 16 (2024).

20. Lichert, F. Interplay between malnutrition and tuberculosis in India. Aktuelle Ernahrungsmedizin 49, 10 (2024).

21. Chandwe, K. et al. Malnutrition enteropathy in Zambian and Zimbabwean children with severe acute malnutrition: A multi-arm randomized phase II trial. Nat Commun 15, 2910 (2024).

22. Vonasek, B.J. et al. Tuberculosis in children with severe acute malnutrition. Expert Rev Respir Med 16, 273–284 (2022).

23. Sinha, P. et al. Undernutrition and Tuberculosis: Public Health Implications. J Infect Dis 219, 1356–1363 (2019).

24. Puri, M., Miranda-Hernandez, S., Subbian, S. & Kupz, A. Repurposing mucosal delivery devices for live attenuated tuberculosis vaccines. Front. Immunol. 14, 1159084 (2023).

25. Dockrell, H.M. & Butkeviciute, E. Can what have we learnt about BCG vaccination in the last 20 years help us to design a better tuberculosis vaccine? Vaccine 40, 1525–1533 (2022).

26. McShane, H. Revaccination with BCG: does it work? The Lancet Infectious Diseases 24, 559–560 (2024).

27. Bekker, L.G. et al. A phase 1b randomized study of the safety and immunological responses to vaccination with H4:IC31, H56:IC31, and BCG revaccination in Mycobacterium tuberculosis-uninfected adolescents in Cape Town, South Africa. EClinicalMedicine 21, 100313 (2020).

28. Singh, M. et al. Efficacy and safety of VPM1002 and Immuvac in preventing tuberculosis: phase 3 randomised clinical trial (PreVenTB trial). BMJ 393, e085716 (2026).

29. StopTBPartnership; Vaccine Pipeline. https://newtbvaccines.org/tb-vaccine-pipeline/

30. Heijmenberg, I. et al. ESX-5-targeted export of ESAT-6 in BCG combines enhanced immunogenicity & efficacy against murine tuberculosis with low virulence and reduced persistence. Vaccine 39, 7265–7276 (2021).

31. Lynn, D.J., Benson, S.C., Lynn, M.A. & Pulendran, B. Modulation of immune responses to vaccination by the microbiota: implications and potential mechanisms. Nat Rev Immunol 22, 33–46 (2022).

32. Shah, T., Shah, Z., Baloch, Z. & Cui, X. The role of microbiota in respiratory health and diseases, particularly in tuberculosis. Biomed Pharmacother 143, 112108 (2021).

33. Eshetie, S. & Van Soolingen, D. The respiratory microbiota: New insights into pulmonary tuberculosis. BMC Infectious Diseases 19 (2019).

34. Budden, K.F. et al. Functional effects of the microbiota in chronic respiratory disease. The Lancet Respiratory Medicine 7, 907–920 (2019).

35. Man, W.H., de Steenhuijsen Piters, W.A.A. & Bogaert, D. The microbiota of the respiratory tract: gatekeeper to respiratory health. Nature Reviews Microbiology 15, 259–270 (2017).

36. Morris, A. et al. Comparison of the respiratory microbiome in healthy nonsmokers and smokers. Am J Respir Crit Care Med 187, 1067–1075 (2013).

37. Khan, N. et al. Intestinal dysbiosis compromises alveolar macrophage immunity to Mycobacterium tuberculosis. Mucosal Immunol. 12, 772–783 (2019).

38. Kennedy, E.A., King, K.Y. & Baldridge, M.T. Mouse Microbiota Models: Comparing Germ-Free Mice and Antibiotics Treatment as Tools for Modifying Gut Bacteria. Front Physiol 9, 1534 (2018).

39. Becattini, S., Taur, Y. & Pamer, E.G. Antibiotic-Induced Changes in the Intestinal Microbiota and Disease. Trends Mol Med 22, 458–478 (2016).

40. Kuypers, M., Despot, T. & Mallevaey, T. Dirty mice join the immunologist’s toolkit. Microbes Infect 23, 104817 (2021).

41. Masopust, D., Sivula, C.P. & Jameson, S.C. Of Mice, Dirty Mice, and Men: Using Mice To Understand Human Immunology. J Immunol 199, 383–388 (2017).

42. Hamilton, S.E. et al. New Insights into the Immune System Using Dirty Mice. J Immunol 205, 3–11 (2020).

43. Choi, Y.H. et al. Safety and Immunogenicity of the ID93 + GLA-SE Tuberculosis Vaccine in BCG-Vaccinated Healthy Adults: A Randomized, Double-Blind, Placebo-Controlled Phase 2 Trial. Infectious Diseases and Therapy 12, 1605–1624 (2023).

44. Cotton, M.F. et al. Safety and immunogenicity of VPM1002 versus BCG in South African newborn babies: a randomised, phase 2 non-inferiority double-blind controlled trial. Lancet Infect Dis 22, 1472–1483 (2022).

45. Dos Santos, P.C.P., et al. Effect of BCG vaccination against Mycobacterium tuberculosis infection in adult Brazilian health-care workers: a nested clinical trial. Lancet Infect Dis 24, 594–601 (2024).

46. Audran, R. et al. Randomised, double-blind, controlled phase 1 trial of the candidate tuberculosis vaccine ChAdOx1-85A delivered by aerosol versus intramuscular route. Journal of Infection 89 (2024).

47. Derrick, S.C., Kolibab, K., Yang, A. & Morris, S.L. Intranasal administration of Mycobacterium bovis BCG induces superior protection against aerosol infection with Mycobacterium tuberculosis in mice. Clin Vaccine Immunol 21, 1443–1451 (2014).

48. Aditama, W., Sitepu, F.Y. & Depari, E. Having contact history with tb active cases and malnutrition as risk factors of TB incidence: A cross-sectional study in North Sumatera, Indonesia. Malaysian Journal of Public Health Medicine 20, 192–198 (2020).

49. Macfarlane, A. Tuberculosis and malnutrition. Br Med J 2, 348 (1947).

50. van Lettow, M. et al. Micronutrient malnutrition and wasting in adults with pulmonary tuberculosis with and without HIV co-infection in Malawi. BMC Infect Dis 4, 61 (2004).

51. Huang, L., Nazarova, E.V., Tan, S., Liu, Y. & Russell, D.G. Growth of Mycobacterium tuberculosis in vivo segregates with host macrophage metabolism and ontogeny. J Exp Med 215, 1135–1152 (2018).

52. Laval, T., Chaumont, L. & Demangel, C. Not too fat to fight: The emerging role of macrophage fatty acid metabolism in immunity to Mycobacterium tuberculosis. Immunol Rev 301, 84–97 (2021).

53. Segal, W. & Bloch, H. Biochemical differentiation of Mycobacterium tuberculosis grown in vivo and in vitro. J. Bacteriol (1956).

54. Lee, W., VanderVen, B.C., Fahey, R.J. & Russell, D.G. Intracellular Mycobacterium tuberculosis exploits host-derived fatty acids to limit metabolic stress. J Biol Chem 288, 6788–6800 (2013).

55. Palma, C. et al. Caloric Restriction Promotes Immunometabolic Reprogramming Leading to Protection from Tuberculosis. Cell Metab 33, 300–318 e312 (2021).

56. Mishra, M. et al. Exoproteome of calorie-restricted humans identifies complement deactivation as an immunometabolic checkpoint reducing inflammaging. Nat Aging 6, 1064–1078 (2026).

57. Craig, W.J. Nutrition Concerns and Health Effects of Vegetarian Diets. Nutrition in Clinical Practice 25, 613–620 (2010).

58. Widmer, R.J., Flammer, A.J., Lerman, L.O. & Lerman, A. The Mediterranean diet, its components, and cardiovascular disease. Am J Med 128, 229–238 (2015).

59. Passos, B.B.S., Araújo-Pereira, M., Vinhaes, C.L., Amaral, E.P. & Andrade, B.B. The role of ESAT-6 in tuberculosis immunopathology. Front. Immunol. 15 (2024).

60. Herp, S. et al. Mucispirillum schaedleri Antagonizes Salmonella Virulence to Protect Mice against Colitis. Cell Host Microbe 25, 681–694 e688 (2019).

61. Zhang, Y. et al. Dubosiella newyorkensis modulates immune tolerance in colitis via the L-lysine-activated AhR-IDO1-Kyn pathway. Nat Commun 15, 1333 (2024).

62. Facchin, S., Calgaro, M. & Savarino, E.V. Rethinking Short-Chain Fatty Acids: A Closer Look at Propionate in Inflammation, Metabolism, and Mucosal Homeostasis. Cells 14 (2025).

63. Ferreira-Paes, T., Seixas-Costa, P. & Almeida-Amaral, E.E. Validation of a Feed Protocol in a Mouse Model That Mimics Marasmic Malnutrition. Front Vet Sci 8, 757136 (2021).

64. Chandrasekaran, P., Saravanan, N., Bethunaickan, R. & Tripathy, S. Malnutrition: Modulator of Immune Responses in Tuberculosis. Front Immunol 8, 1316 (2017).

65. Levy, M. et al. Microbiota-Modulated Metabolites Shape the Intestinal Microenvironment by Regulating NLRP6 Inflammasome Signaling. Cell 163, 1428–1443 (2015).

66. Bolyen, E. et al. Reproducible, interactive, scalable and extensible microbiome data science using QIIME 2. Nat Biotechnol 37, 852–857 (2019).

67. Callahan, B.J. et al. DADA2: High-resolution sample inference from Illumina amplicon data. Nat Methods 13, 581–583 (2016).

68. Parks, D.H. et al. GTDB: an ongoing census of bacterial and archaeal diversity through a phylogenetically consistent, rank normalized and complete genome-based taxonomy. Nucleic Acids Res 50, D785–D794 (2022).

69. Quast, C. et al. The SILVA ribosomal RNA gene database project: improved data processing and web-based tools. Nucleic Acids Res 41, D590–596 (2013).

70. Cole, J.R. et al. The ribosomal database project (RDP-II): introducing myRDP space and quality controlled public data. Nucleic Acids Res 35, D169–172 (2007).

71. Kumar, M. & Field, M.A. (2025). biorxiv. 10.1101/2025.11.03.686403

72. Team, R.C. A language and environment for statistical computing. Foundation for Statistical Computing, Vienna, Austria. (2013).

73. McMurdie, P.J. & Holmes, S. phyloseq: an R package for reproducible interactive analysis and graphics of microbiome census data. PLoS One 8, e61217 (2013).

74. H, W., F.R., L, H., K, M. & D, V. dplyr: A Grammar of Data Manipulation. (2025).

75. Wickam, H. ggplot2: Elegant Graphics for Data Analysis. Springer-Verlag New York. (2016).

