## Supplementary Materials for "Malnourishment and expanded microbiome maintain superior efficacy of the recombinant tuberculosis vaccine BCG::ESAT-6-PE25SS"

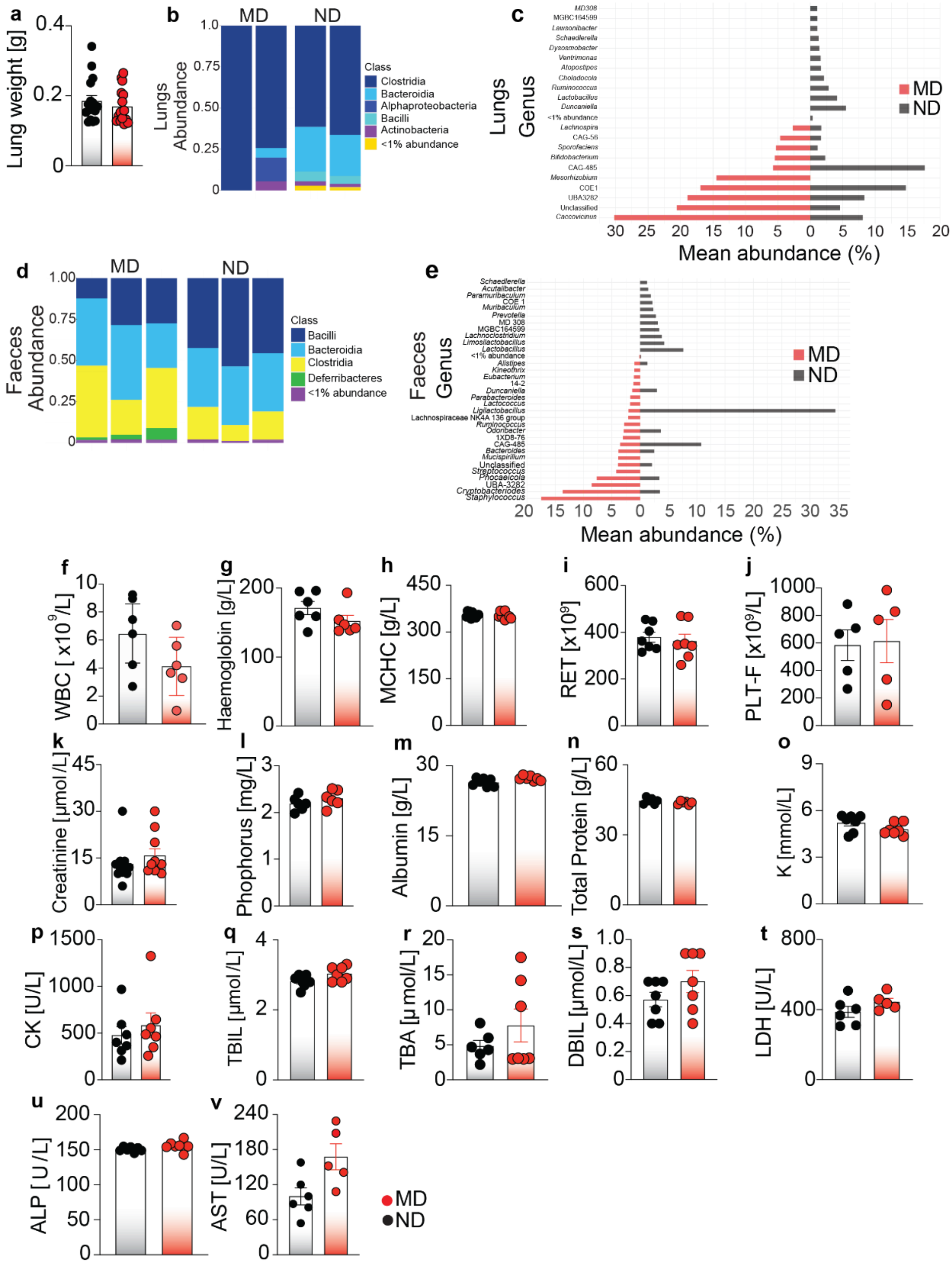


**Supplementary Figure 1: a)** Lung weight of MD- and ND-fed mice at 14 days after diet intervention. **b, d)** Relative abundance of bacterial classes in lungs (**b**) and faeces (**d**) in ND- and MD-fed diet **c-e)** Bidirectional plots of bacterial genera in lungs (**c**) and faeces (**e**) of ND-fed and MD-fed diets **f-v)** White blood cell count (**f**), plasma levels of hemoglobulin (**g**), mean corpuscular hemoglobulin concentration (MCHC) (**h**), reticulocytes (RET; (**i**), platelets (PLT-F (**j**), creatinine (**k**), phosphorus (**l**), albumin (**m**), total protein (**n**), potassium (**o**), creatine kinase (CK) (**p**), total Bilirubin (TBIL) (**q**), total bile acid (TBA) (**r**), direct bilirubin (DBIL) (**s**), lactate dehydrogenase (**t**), alkaline phosphatase (ALP) (**u**), aspartate aminotransferase (AST) (**v**). Results are representative of five pooled independent experiments n=5-16 mice per group. Statistical significance was calculated using Mann-Whitney test. MD, malnourished diet; ND, normal diet.

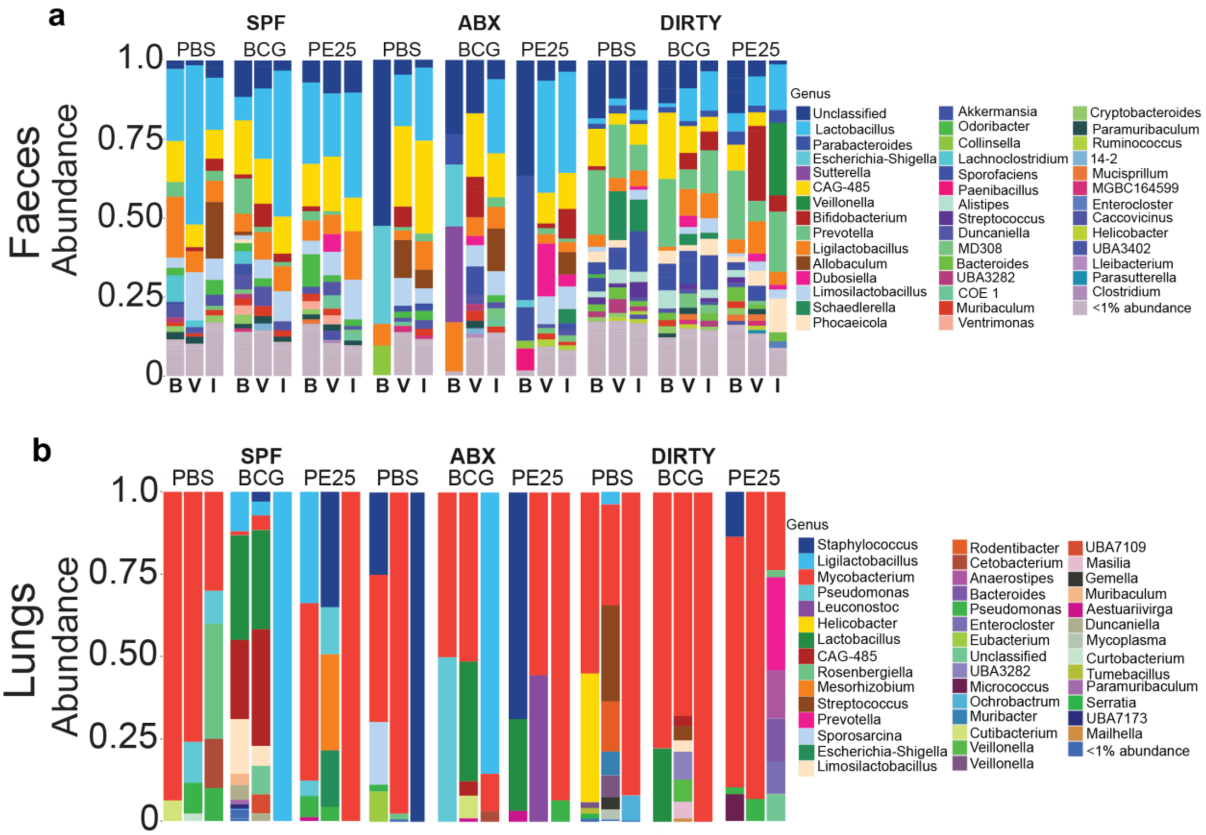


**Supplementary Figure 2: a-b)** Relative abundance of bacterial genera in faeces (**a**) and lungs (**b**) of SPF, AXB and dirty mice. B, baseline; V, 60 days after vaccination; I, 45 days after *Mtb* infection

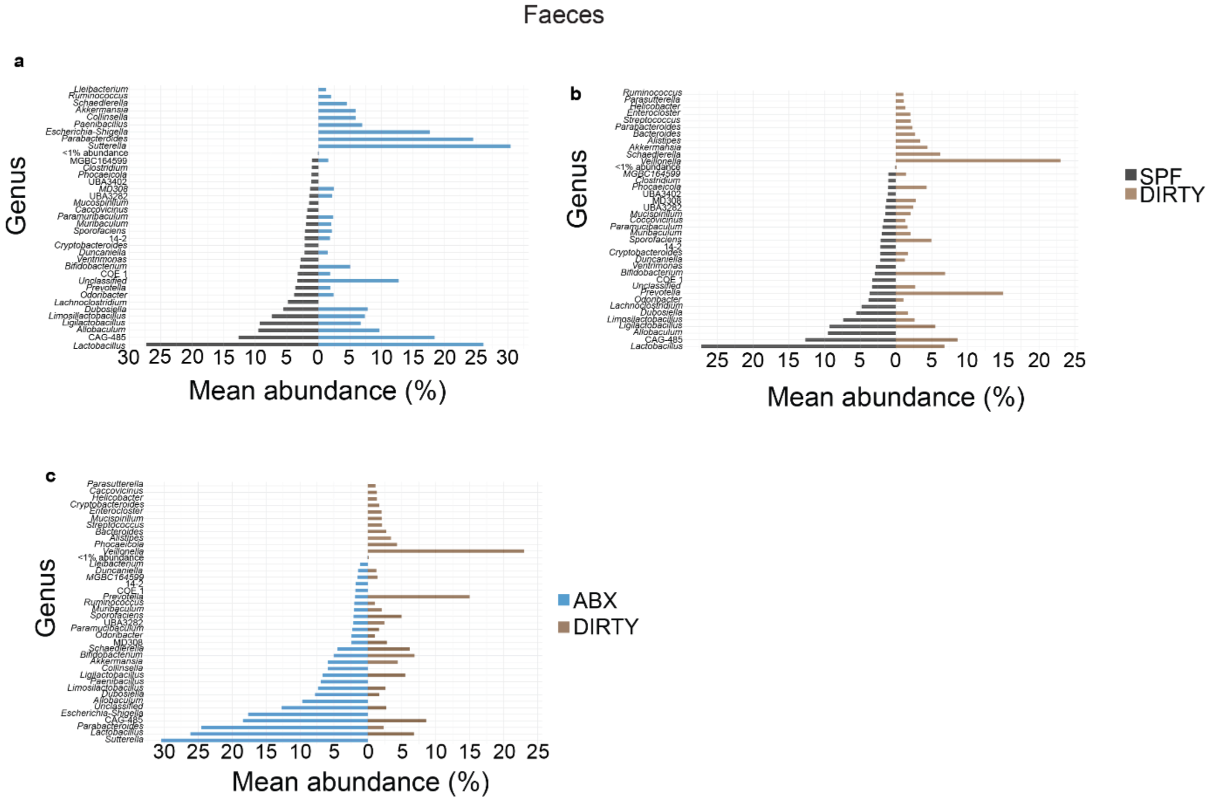


**Supplementary Figure 3**: **a-c**) Bidirectional abundance plot from faeces comparing SPF and ABX mice (**a**), SPF and dirty mice (**b**) and ABX and dirty mice (**c**).

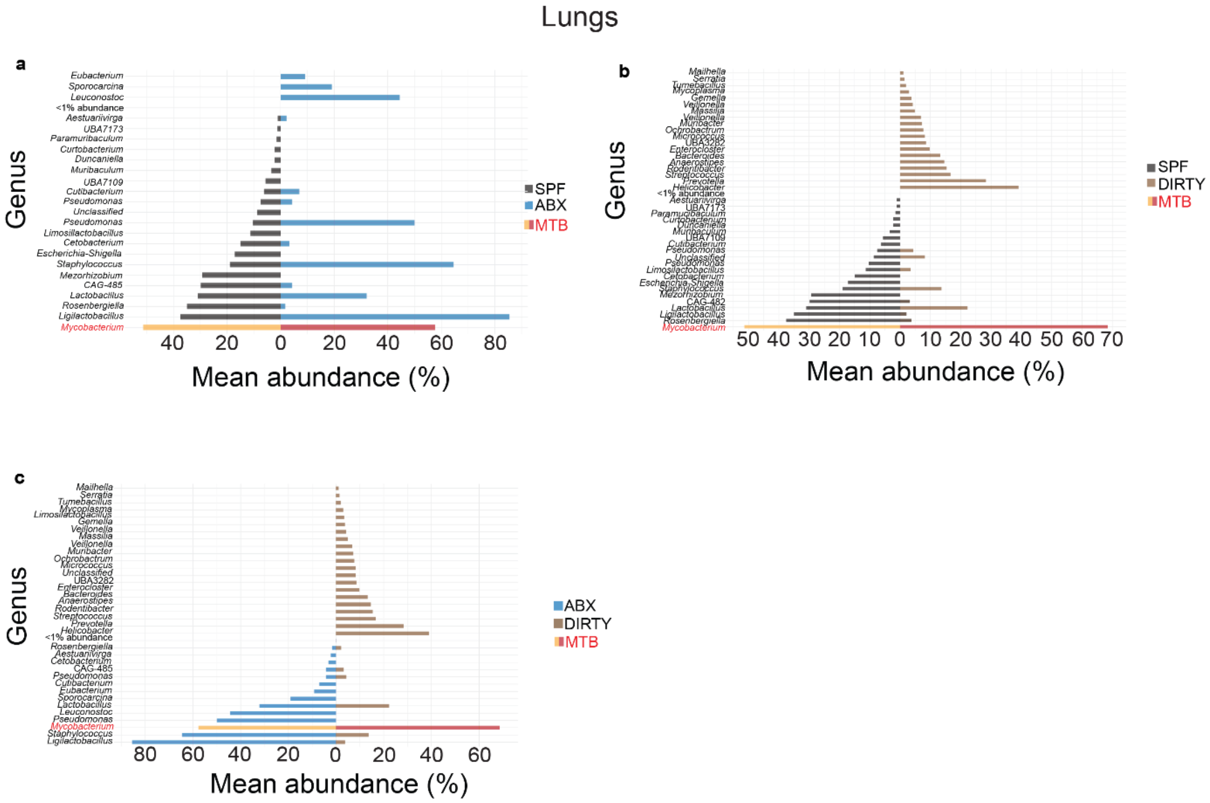


**Supplementary Figure 4: a-c**) Bidirectional abundance plot from lungs comparing SPF and ABX mice (**a**), SPF and dirty mice (**b**) and ABX and dirty mice (**c**).

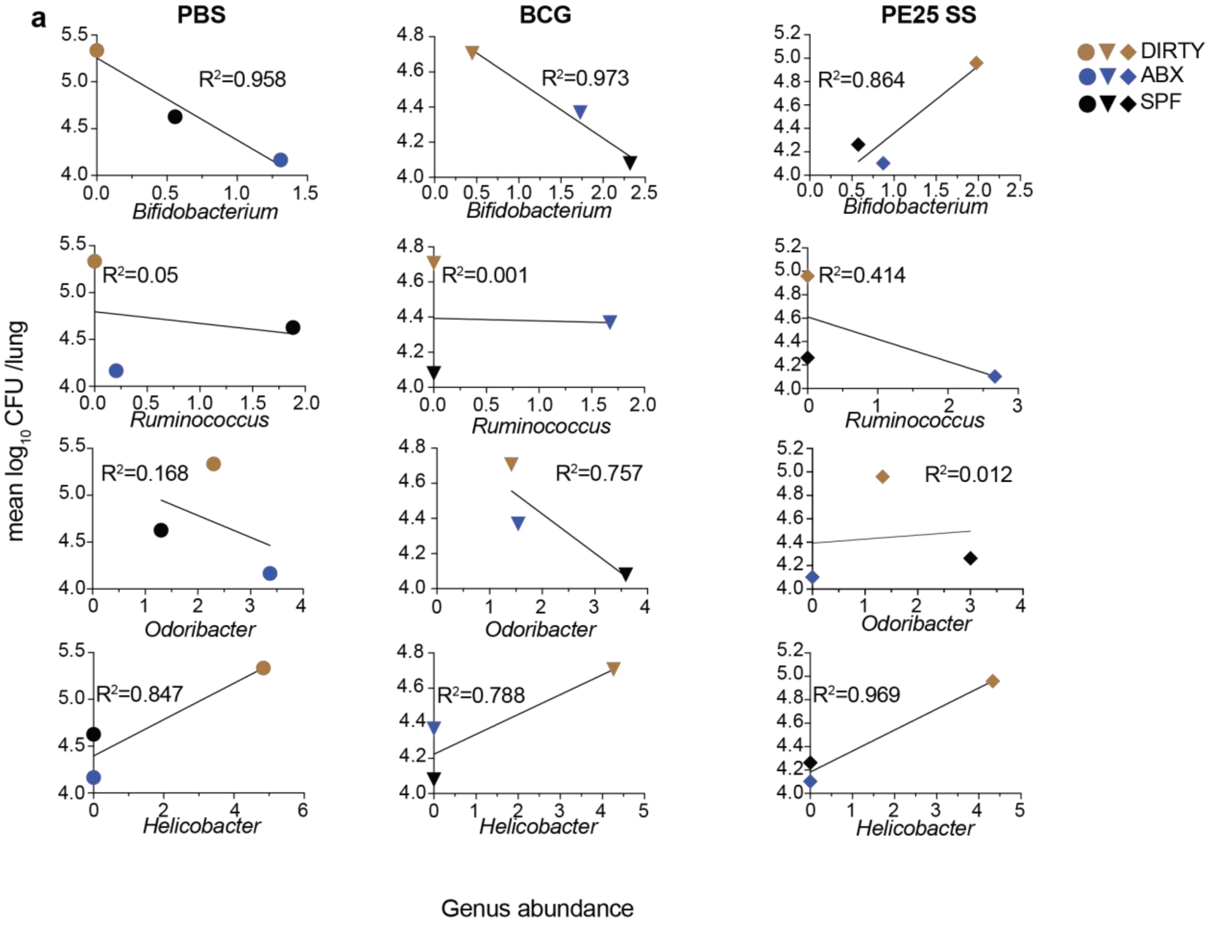

a
PBS
BCG
PE25 SS
DIRTY
ABX
SPF
R²=0.958
R²=0.973
R²=0.864
R²=0.05
R²=0.001
R²=0.414
R²=0.168
R²=0.757
R²=0.012
R²=0.847
R²=0.788
R²=0.969
Bifidobacterium
Ruminococcus
Odoribacter
Helicobacter
mean log10 CFU /lung
Genus abundance

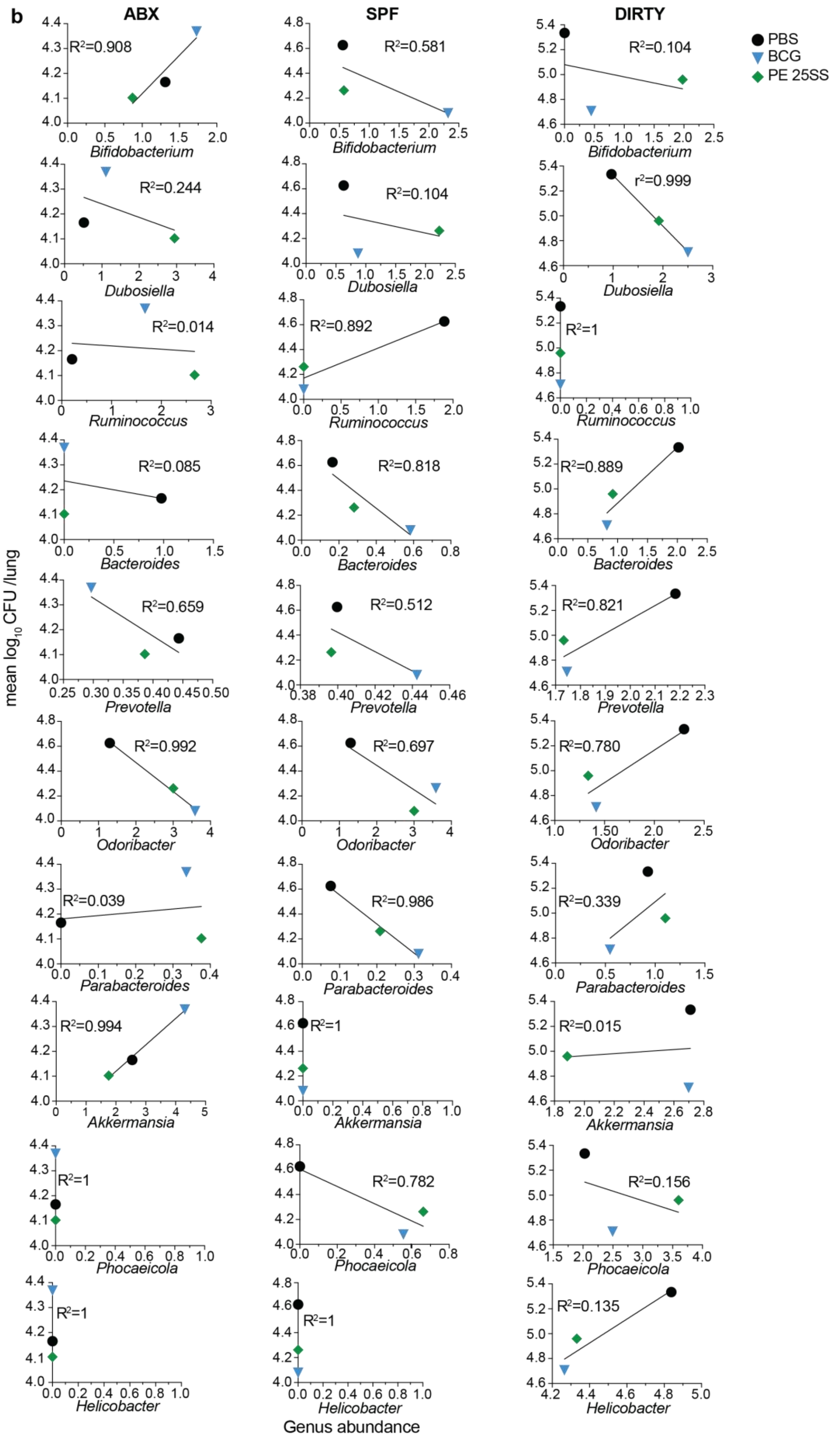
b
ABX
SPF
DIRTY
PBS
BCG
PE 25SS
mean log10 CFU /lung
Genus abundance
Bifidobacterium
Dubosiella
Ruminococcus
Bacteroides
Prevotella
Odoribacter
Parabacteroides
Akkermansia
Phocaeicola
Helicobacter
R²=0.908
R²=0.581
R²=0.104
R²=0.244
R²=0.104
r²=0.999
R²=0.014
R²=0.892
R²=1
R²=0.085
R²=0.818
R²=0.889
R²=0.659
R²=0.512
R²=0.821
R²=0.992
R²=0.697
R²=0.780
R²=0.039
R²=0.986
R²=0.339
R²=0.994
R²=1
R²=0.015
R²=1
R²=0.782
R²=0.156
R²=1
R²=1
R²=0.135

**Supplementary Figure 5: Correlation analysis between lung *Mtb* burden and bacterial abundance. a)** Mean lung *Mtb* CFU versus bacterial abundance plots, including $R^2$ values for selected bacterial genera (*Bifidobacterium, Ruminococcus, Odoribacter and Helicobacter)* in unvaccinated, BCG-vaccinated and BCG::ESAT-6-PE25SS-vaccinated groups. **b**) Mean lung *Mtb* CFU versus bacterial abundance plots, including $R^2$ values for selected bacterial genera in SPF, ABX and dirty mice.
